# Evaluating Methods for the Prediction of Cell Type-Specific Enhancers in the Mammalian Cortex

**DOI:** 10.1101/2024.08.21.609075

**Authors:** Nelson J. Johansen, Niklas Kempynck, Nathan R. Zemke, Saroja Somasundaram, Seppe De Winter, Marcus Hooper, Deepanjali Dwivedi, Ruchi Lohia, Fabien Wehbe, Bocheng Li, Darina Abaffyová, Ethan J. Armand, Julie De Man, Eren Can Eksi, Nikolai Hecker, Gert Hulselmans, Vasilis Konstantakos, David Mauduit, John K. Mich, Gabriele Partel, Tanya L. Daigle, Boaz P. Levi, Kai Zhang, Yoshiaki Tanaka, Jesse Gillis, Jonathan T. Ting, Yoav Ben-Simon, Jeremy Miller, Joseph R. Ecker, Bing Ren, Stein Aerts, Ed S. Lein, Bosiljka Tasic, Trygve E. Bakken

## Abstract

Identifying cell type-specific enhancers in the brain is critical to building genetic tools for investigating the mammalian brain. Computational methods for functional enhancer prediction have been proposed and validated in the fruit fly and not yet the mammalian brain. We organized the ‘Brain Initiative Cell Census Network (BICCN) Challenge: Predicting Functional Cell Type-Specific Enhancers from Cross-Species Multi-Omics’ to assess machine learning and feature-based methods designed to nominate enhancer DNA sequences to target cell types in the mouse cortex. Methods were evaluated based on *in vivo* validation data from hundreds of cortical cell type-specific enhancers that were previously packaged into individual AAV vectors and retro-orbitally injected into mice. We find that open chromatin was a key predictor of functional enhancers, and sequence models improved prediction of non-functional enhancers that can be deprioritized as opposed to pursued for *in vivo* testing. Sequence models also identified cell type-specific transcription factor codes that can guide designs of *in silico* enhancers. This community challenge establishes a benchmark for enhancer prioritization algorithms and reveals computational approaches and molecular information that are crucial for identifying functional enhancers in mammalian cortical cell types. The results of this challenge bring us closer to understanding the complex gene regulatory landscape of the mammalian cortex and to designing more efficient genetic tools to target cortical cell types.

## Introduction

The mammalian neocortex, responsible for higher-order cognitive functions and sensorimotor processing, includes the primary motor cortex (M1), which facilitates fine motor movement and is composed of diverse cell types with distinct molecular signatures ^1,2^. Some neurodegenerative diseases, including Parkinson’s, Huntington’s, and amyotrophic lateral sclerosis (ALS), affect specific M1 cell types and result in impaired coordination and dexterity ^3^. There is an urgent need for genetic tools to selectively access vulnerable cell populations to probe cortical circuit function and treat disease. To this end, cell type-selective candidate enhancers have been identified based on single cell genomic profiling of mouse cortex and have been used to create recombinant adeno-associated virus (AAV) vectors to drive exogenous transgene expression in the predicted target cell types ^4–6^. With increasingly comprehensive molecular phenotyping of the brain ^7–11^, there is the prospect of developing AAV tools to target a wide diversity of cell types across the brain.

Identification of cell type-specific viral tools remains challenging because experimental validation is low-throughput and expensive. A recent study by Ben-Simon et al. 2024 ^6^ tested 825 enhancers selected based on specificity of open chromatin within a cortical cell type of interest and achieved an overall average success rate of 30%. Consequently, the field needs new computational approaches to improve functional enhancer prediction ^9^ and accelerate viral tool development that integrates comprehensive molecular profiling within and across species. However, it remains poorly understood how well different approaches predict functional over non-functional enhancer genomic sequences. Community-guided challenges have demonstrated their effectiveness in rigorously evaluating novel computational methods and advancing knowledge in the genomics field ^12^. To date, there has been no community-led effort to address the prioritization of cell type-specific enhancers using cutting edge cell type-resolved atlases from multiple species.

In this study, we present the BICCN Challenge, where six teams from computational biology labs across the world participated to predict functional cortical cell type-specific enhancers. We introduce a community-driven benchmark and metrics to identify top-performing approaches that prioritize functional, cell type-specific viral tools. By investigating the computational approaches and biological priors used by high-performance methods, we aim to contribute to the refinement of functional enhancer prediction to selectively target cell types in the mammalian cortex.

## Results

### A community challenge to predict functional enhancers

We provided teams with a comparative multi-omics study of M1 by Zemke et al. ^13^ that measured the molecular profiles of individual nuclei using single-cell multi-omics and single-cell methyl-Hi-C (snm3C) in human, macaque, marmoset and mouse (**Fig. 1A**). Multiple species were included because molecular and DNA sequence patterns associated with enhancer activity are conserved in the mammalian brain ^1,14^. Teams were asked to combine cross-species genomics measurements and biological priors to prioritize cell type-specific and functional enhancer elements (**Fig. 1B**). We asked teams to provide the top 10,000 putative enhancers for each cell type to increase the likelihood of validated enhancers being included in the rankings.

**Figure 1.**
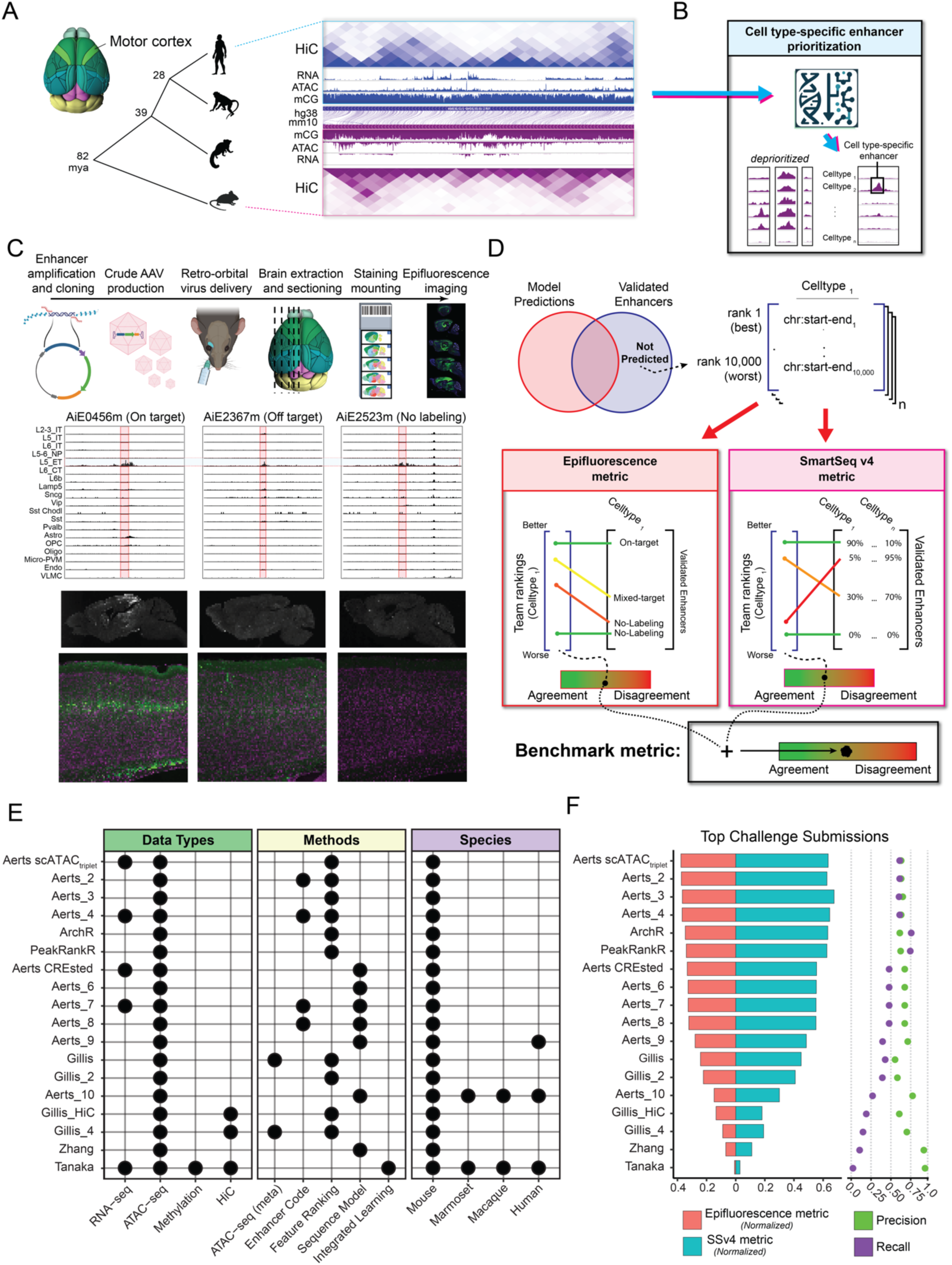
Overview of the enhancer prioritization challenge. (A) Single nucleus multi-omic data from primary motor cortex of human, macaque, marmoset, and mouse. mya, million years ago. (B) Schematic of the computational challenge to prioritize candidate cell type-specific enhancers. (C) Overview of AAV construction, cell type ATAC-seq specificity, and screening of *in vivo* activity in the mouse brain for three candidate L5 ET enhancers. (D) Teams predicted and ranked 10,000 candidate enhancers for each of 19 cortical cell types and were scored based on prioritization of strong, On-Target enhancers. (E) Combinations of data and methods for top team submissions. (F) Normalized benchmark metrics (**Methods**) based on epifluorescence and SSv4 from *in vivo* screening.

Teams’ predictions were evaluated against 677 recombinant adeno-associated virus (AAV) vectors among 825 vectors that were designed to label 19 M1 cell subclasses and assessed for *in vivo* enhancer specificity and brightness in the mouse brain ^6^. Validation data for 148 enhancers were released after the challenge and were therefore excluded from the analysis. The cell type-specificity in the neocortex of each validated enhancer virus was assessed through manual evaluation of epifluorescence images, following the methodology outlined in Ben-Simon et al. ^6^. The enhancers were classified into four groups: (1) On-Target (N=202), specific labeling of the targeted cell type; (2) Off-Target (N=96), labeling of non-targeted cell types in the neocortex; (3) Mixed-Target (N=100), labeling of targeted and non-targeted cell type(s); and (4) No-Labeling (N=279), no fluorescence in the mouse neocortex (**Fig. 1C, Supplemental Table 1**). To further quantify the in vivo activity of 191 enhancers, SYFP2-positive cells were extracted from the primary visual cortex (V1) and analyzed with Smart-seq v4 (SSv4) sequencing^6^.

The challenge was run over a 3-month period. Participants submitted enhancer lists at several intervals, and performance was reported on a public leaderboard^15^. In the final evaluation round, teams provided a detailed description of their approach (**Methods**). To rigorously evaluate methods, we developed a benchmark metric that is optimized when enhancers with On-Target activity are ranked highest, and Mixed-Target, Off-Target and No-Labeling enhancers are ranked lowest for the respective cell type (**Fig. 1D, Methods**). Final scores per method were computed based on both epifluorescence and SSv4 metrics using the entire corpus of validated enhancers. The benchmark metric and an automated scoring tool are hosted via GitHub^15^ for the community to fairly evaluate novel enhancer prioritization methods.

### Top performing submissions focused on ATAC-seq specificity

Challenge participants from 5 teams contributed 79 submissions that comprised 16 unique enhancer prioritization methods, and the ArchR ^16^ method was included as a performance baseline due to its popularity as a single cell ATAC-seq processing pipeline. We grouped the methods into five broad categories based on the included enhancer features: (1) ATAC-seq (meta), (2) Enhancer codes, (3) Feature Ranking, (4) Sequence Model and (5) Integration Model (**Fig. 1E**). The top three teams (Aerts, ArchR and PeakRankR) achieved comparable performance with normalized scores between 0.36 and 0.41, while the remaining teams achieved scores below 0.27 (**Fig. 1F**). Comparing methods based on precision and recall identifies that even top performing teams are only able to achieve moderate accuracy in recovering On-Target enhancers (F1-score, Aerts: 0.41, ArchR: 0.37, PeakRankR: 0.36) Teams submitted diverse approaches and markedly improved during the challenge (**Fig. 1E-F**, **Supplemental Fig. 1**, **Supplement Table 2**). The final challenge ranks were statistically robust (P < 0.05) for 55 of 79 submissions based on a bootstrapping analysis (**Supplemental Fig. 2, Supplemental Table 2**).

High-performing teams used similar approaches that leveraged ATAC-seq features, including differential chromatin accessibility (specificity) and signal strength at a given enhancer genomic location. The top performing submission gained a slight advantage by including RNA-seq and leveraging SCENIC+ ^17^ to predict cell type-specific transcription factor (TF)-enhancer-gene triplets. The runner-up baseline method, ArchR, used careful selection of background cells to minimize biases such as transcription start site (TSS) enrichment when performing pairwise statistical tests. The third-ranking team, PeakRankR, calculated three ATAC-seq metrics – specificity, magnitude and coverage – that were combined to discriminate cell type-selective enhancers.

Comparison of team submissions highlighted the genomics data, biological priors, and methodology that enabled accurate prediction of functional cell type enhancers (**Supplemental Fig. 3, Supplemental Table 2**). While all submissions used mouse ATAC-seq data, they varied in how the signal was normalized to handle batch effects and how cell type specificity was calculated. Surprisingly, inclusion of DNA-methylation, chromatin folding (HiC), or primate data decreased performance, potentially due to increased model complexity and overfitting. Also, the Tanaka team used all data types but retained only the highest magnitude open chromatin features that left primarily promoters, not enhancers, for the method to prioritize (**Supplemental Text**). Notably, a high performance, runner-up method (Aerts CREsted) employed deep learning models, trained with the CREsted package ^18^, that learned to predict open chromatin from DNA sequence. Sequence models are of particular interest since they have the potential to identify TF motifs that make up cell type-specific enhancer codes ^19,20^, including repressive elements.

### Meta-analysis of performance across teams and cell types

To assess whether teams identified similar or distinct enhancers, we computed pairwise intersections of submissions from the top-performing methods (**Fig. 2A**). Methods with similar performance predicted similar enhancers, and the top three methods (Aerts scATAC_triplet_, ArchR and PeakRankR) had the highest agreement. Many On-Target enhancers were identified across several approaches (**Fig. 2B**), yet some were not recovered by any method, likely due to low ATAC-seq signal (**Supplemental Fig. 4**).

**Figure 2.**
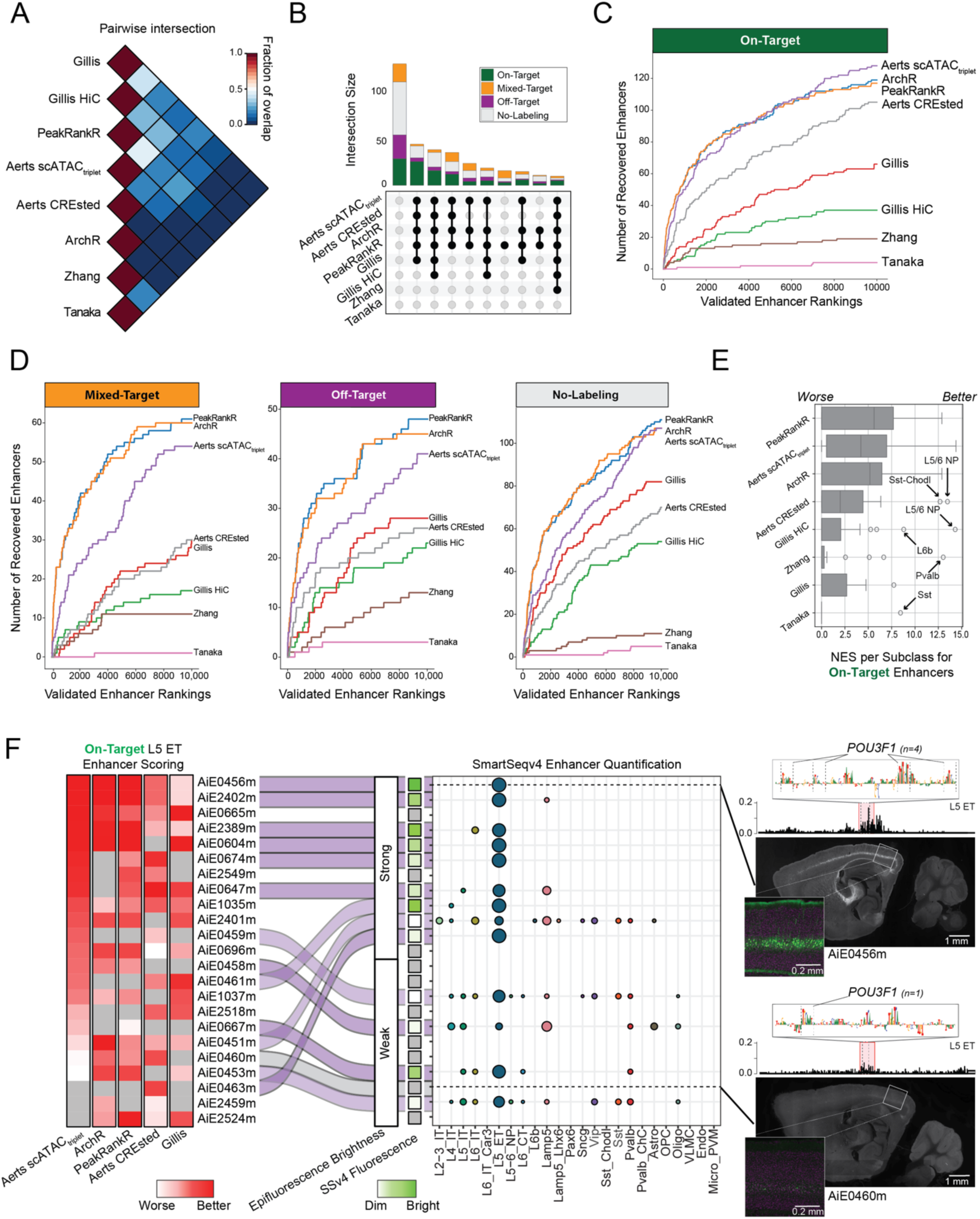
Comparison of team enhancer rankings. (A) Average proportion of ranked enhancers that overlap between pairs of team submissions for all cell types. (B) Upset plot showing the number of validated enhancers that were identified by sets of submissions (C,D) Rates of identification of (C) On-Target and (D) Mixed-Target, Off-Target and No-Labeling enhancers. (E) Comparison of methods based on distributions of normalized enrichment scores (NES). For each method and cell type specific ranking, NES measures the area under the recovery curve (AUC) up to the 1,000^th^ element compared to a random ranking. (F) Heatmap ordered by Aerts scATAC_triplet_ scoring of L5 ET enhancers and summary of validation results. Examples of a strong (AiE0456m) and weak (AiE0460m) enhancer with *Pou3f1* motifs identified by the CREsted model in the highlighted region. AiE0456m was also validated with SSv4.

The Aerts, PeakRankR, and ArchR methods consistently included On-Target enhancers in the top rankings (**Fig. 2C**), while the Aerts method deprioritized more Mixed-Target, Off-Target and No-Labeling (**Fig. 2D**). From these recovery curves, we calculated the normalized enrichment score (NES) that quantifies enrichment of functional enhancers at the top of a method’s ranking (**Fig. 2E**). Interestingly, compared to the top-performing methods, methods that used DNA sequence (Aerts CREsted) and HiC (Gillis) data better prioritized enhancers for L6b neurons, and the CREsted method better prioritized enhancers for low-abundance Sst Chodl inhibitory neurons (**Fig. 2E**, **Supplemental Figs. 5-6, Supplemental Table 2**). In addition, Aerts scATAC_triplet_ and Aerts CREsted were better than ArchR and PeakRankR at deprioritizing lower quality enhancers for most cell types (**Supplemental Figs. 7-8**).

Next, we compared performance in prioritizing L5 ET enhancers, the largest set of validated enhancers. The top five strongest and most specific On-Target L5 ET enhancers were ranked highly by most methods, with somewhat lower rankings from Gillis (**Fig. 2F**). Most methods correctly scored a strong L5 ET enhancer AiE0456m higher than a weak enhancer AiE0460m, likely due to higher ATAC-seq signal in AiE0456m and the presence of multiple POU3F1 motifs (**Fig. 2F**), a canonical TF of L5 ET neurons ^1^. The strong enhancer AiE0463m had low ATAC-seq signal but multiple POU3F1 motifs and was correctly prioritized only by the Aerts CREsted model (**Fig. 2F**, **Supplemental Fig. 9**). Thus, methods that can learn from DNA sequence have an advantage to prioritize strong and specific enhancers that are associated with chromatin regions with limited accessibility.

### Additional genomic features help predict enhancer activity

Since chromatin accessibility sometimes failed to predict cell type-specific activity, we tested if prediction accuracy could be improved by including additional enhancer features that were not provided to teams in the challenge. These features included (1) H3K27ac, a histone modification found at active enhancers ^21^ from mouse cortical Paired-Tag data ^13,22^, (2) Activity-By-Contact (ABC) scores, the product of chromatin accessibility and contact frequency from HiC ^13,23^, and (3) conservation of chromatin accessibility between human and mouse ^13^. Like chromatin accessibility, cell type-specific H3K27ac and ABC scores were significantly higher for On-Target validated enhancers compared to Mixed-Target, Off-Target, No-Labeling and a random control (**Fig. 3A**). On-Target enhancers had significantly higher conservation of accessibility (**Fig. 3A**) and sequence (**Supplemental Fig. 10**) compared with No-Labeling and Random but not Mixed-Target or Off-Target enhancers. Thus, conservation of accessibility predicted overall enhancer activity, while H3K27ac and ABC predicted cell type-specific enhancer activity.

**Figure 3.**
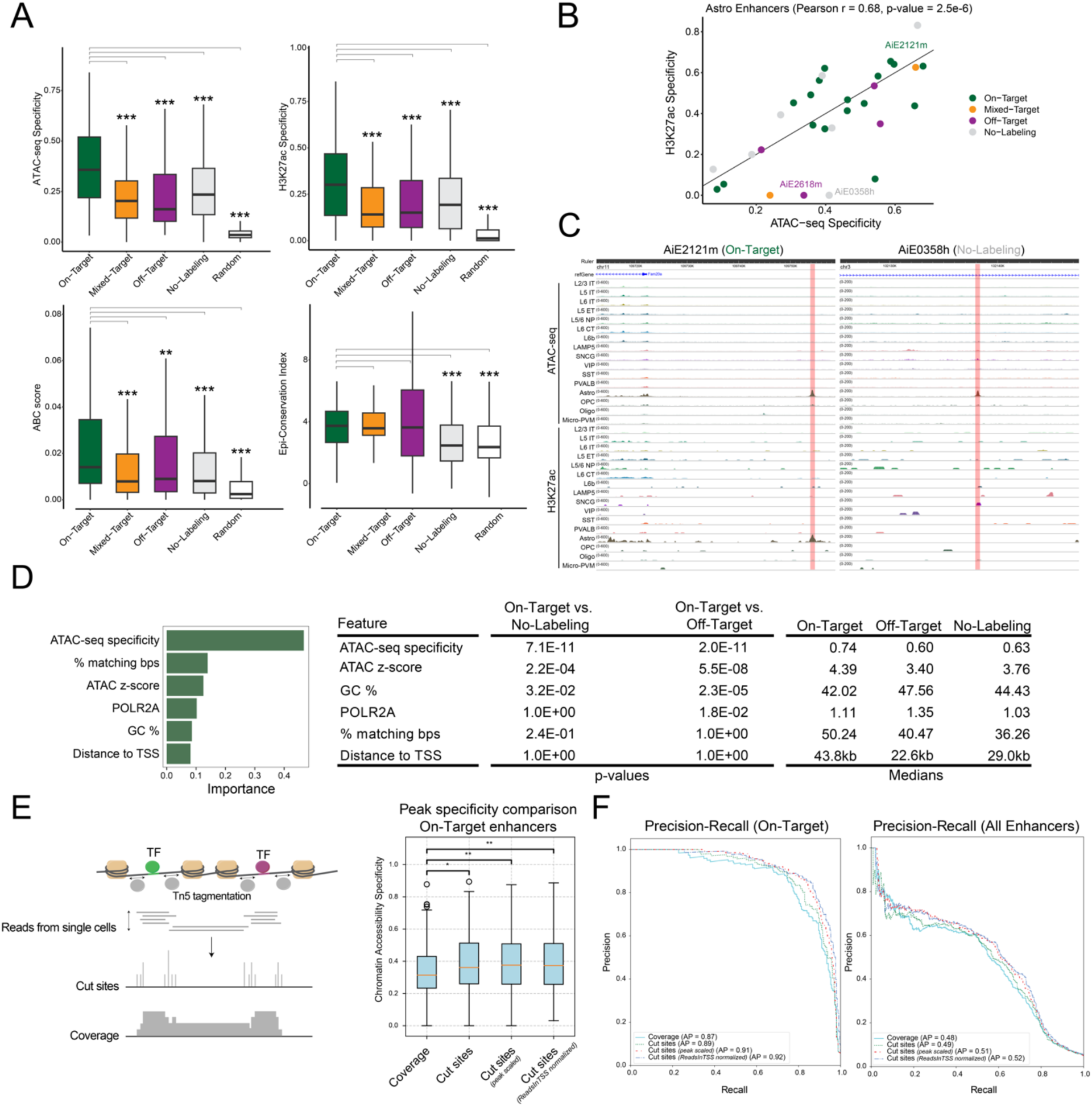
Enhancer features predictive of functional activity. (A) Comparison of molecular features between On-Target and other enhancer categories. ** P < 0.001, *** P < 0.0001 Wilcoxon rank-sum test two-sided, unpaired. (B) Correlation of H3K27ac and ATAC-seq specificity for astrocyte enhancers. (C) Examples of astrocyte enhancers with *in vivo* activity that is better predicted by H3K27ac than ATAC-seq signal. (D) Summary of informative features from a Random Forest model predicting enhancer activity. ANOVA with Tukey post hoc tests, Bonferroni-corrected P-values. (E) Schematic of ATAC-seq peak quantification based on cut sites or coverage. Box plot comparison of peak specificity for all on-target enhancers between different preprocessing methods. Adjusted p-values were obtained through t-tests, *P < 0.05, **P < 0.01. (F) Overall enhancer activity prediction performance from peak specificity for the different methods for On-target (left) and all, except Off-target, (right) enhancers.

For validated enhancers, cell type-specific chromatin accessibility was notably correlated with cell type-specific H3K27ac (r = 0.57) and ABC scores (r = 0.38), but not epigenetic conservation (r = 0.07) (**Fig. 3B, Supplemental Fig. 11**). For example, On-Target enhancer AiE2121m had high chromatin accessibility and H3K27ac specifically in astrocytes (**Fig. 3B,C**). In contrast, No-Labeling element AiE0358h had high chromatin accessibility but not H3K27ac specifically in astrocytes (**Fig. 3B,C**). These examples demonstrate a potential for H3K27ac signal to improve the accuracy of enhancer predictions by distinguishing functional from non-functional chromatin accessible candidate enhancers. Additionally, we trained a binary random forest classification model to predict On-Target enhancers from all enhancers that were Off-target or had no labeling. This approach made use of labelled data and thus is a different approach than the BICCN Challenge submissions described in this manuscript. In the held-out test set, this model resulted in an AUC of 0.75. This model is better able to identify enhancers that are Off-Target, or have no labeling (precision 0.77, recall 0.80) than On-Target enhancers (precision 0.52, recall 0.47). While the precision is relatively low, this model out-performed the initial enhancer selection where only 30% of screened enhancers were scored as On-Target^6^. The most informative features can be leveraged in future enhancer prediction models, including open chromatin metrics (ATAC-seq strength or z-score and cell type specificity), linear sequence conservation (% matching bps) and GC content (**Fig. 3D, Supplemental Fig. 12**).

Finally, we compared differences in ATAC-seq preprocessing methods, as there was some variability in data processing for the top performing differential accessibility rankings. We compared counts-per-million (CPM) normalized coverage and cut site tracks (**Supplemental Fig. 13**) and found that cell type-specific peak heights based on accumulation of the signal inside a peak are significantly (P_adj_ < 0.05) more specific in On-Target enhancers if they are obtained from cut site tracks (**Fig. 3E**). Moreover, additional normalization methods to scale peak heights across cell types (see Methods) further increase specificity. Using specificity as a metric to score enhancer activity in a multilabel classification setting (**Fig. 3F**), we confirmed the increased performance of cut sites-based peaks compared to coverage peaks both for On-Target and all, except Off-Target, validated enhancers. These results demonstrate the benefit of peak normalization, which ensures better comparability of peak heights across cell types.

### Re-scored enhancers validate model predictions

To further investigate differences in predicted versus *in vivo* activity for the 677 tested enhancers, we manually inspected the scATAC-seq signals from Aerts scATAC_triplet_ and prediction and nucleotide contribution scores from Aerts CREsted. These top-performing models used the same CPM-normalized ATAC-seq coverage tracks. We classified each enhancer into one of five categories: ‘Explainable positives’ (48%) have *in vivo* activity and high ATAC specificity and CREsted scores in the corresponding cell type; ‘Explainable negatives’ (25.4%) have no *in vivo* activity and low ATAC or CREsted specificity; ‘Unexplainable positives’ (6.2%) have activity but low ATAC and CREsted specificity; ‘Unexplainable negatives’ (13.9%) have no reported activity yet high ATAC and CREsted scores; and ‘Missing data’ (6.5%) for the remaining enhancers. We visualized enhancers based on the peak-scaled ATAC signal and labeled by the targeted cell type (**Fig. 4A**). Explainable On-Target enhancers were well segregated and overlapped with some unexplainable No-Labeling (negative) enhancers. This suggested that some negative enhancers may have weak *in vivo* activity that was missed in the initial evaluation.

**Figure 4.**
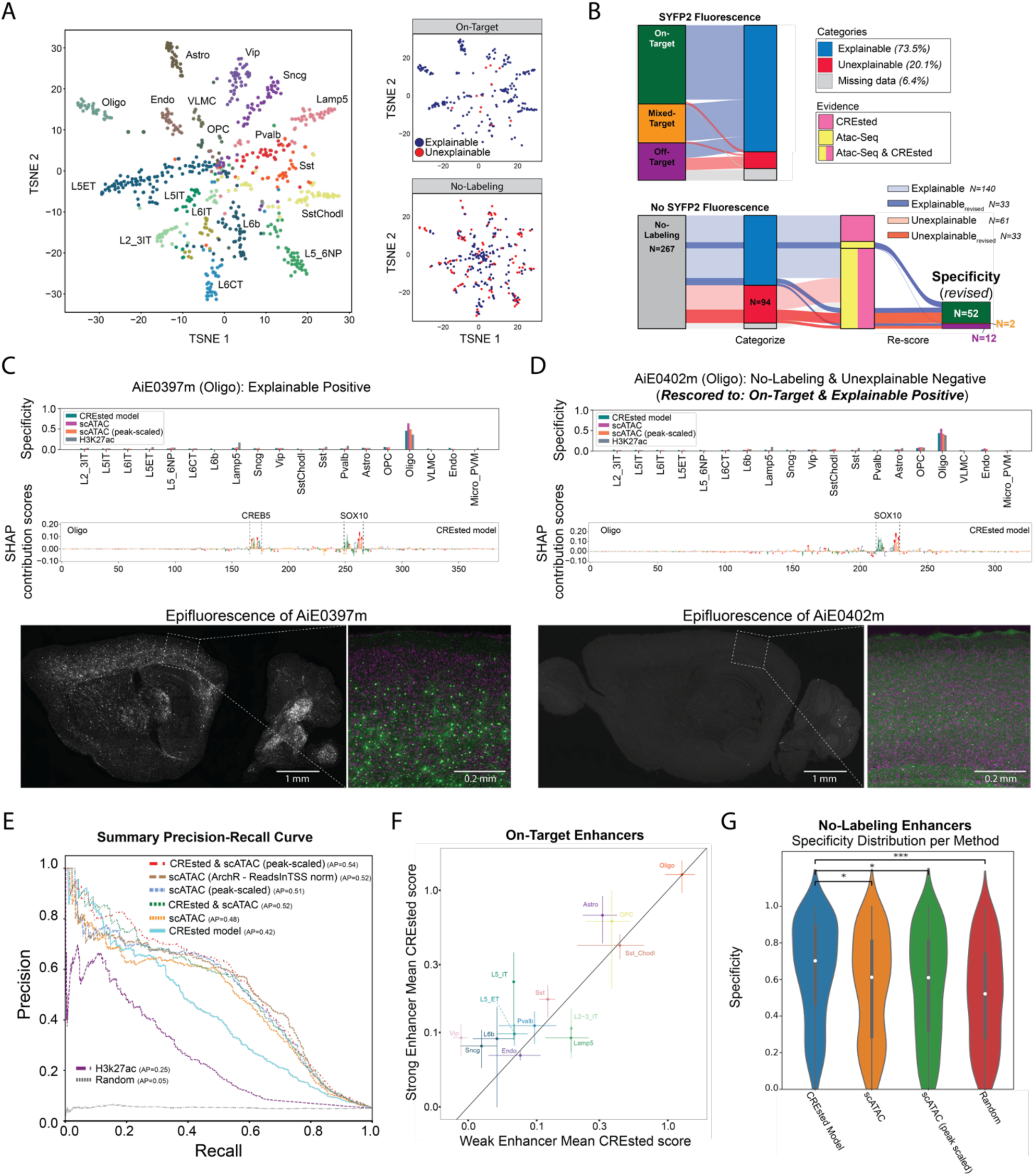
Refinement of models and enhancer screening results. (A) tSNE plots of enhancers based on ATAC-seq specificity and labeled by the targeted cell type. On-Target and No-Labeling enhancers had explainable or unexplainable cell type labeling patterns based on ATAC-seq and DNA sequence (CREsted) model predictions. (B) River plots of enhancer activity, predictions and rescoring of experimental validation data (C-D) Model scores, predicted TF motifs and SYFP fluorescence for two Oligo enhancers with epifluorescence strengths (C) strong On-Target and (D) No-Labeling rescored to weak On-Target activity. (E) Performance of enhancer ranking methods using the rescored enhancer activities. AP, average precision. scATAC included two normalizations: count-normalized coverage pseudobulk or peak-scaled. (F) CREsted model scores for strong and weak On-Target enhancers grouped by cell type. Mean +/-SEM. (G) Comparison of models at identifying No-Labeling enhancers. * P < 0.05, *** P < 0.001, Wilcoxon rank-sum test, Bonferroni-corrected P-values.

We re-evaluated validation data from all 267 No-Labeling enhancers and found that 66 enhancers weakly drove SYFP2 expression in the brain (**Fig. 4B**). 52 of 66 enhancers had weak expression in the targeted cell type and were rescored to On-Target. These enhancers were enriched for unexplainable versus explainable negatives, which supports our predictions of enhancer activity. Interestingly, among the 140 explainable negatives that were confirmed to have no activity after rescoring, 45.7% had ATAC-seq signal in their targeted cell type but no support from the CREsted model. Conversely, only two enhancers (1.3%) contained a specific CREsted prediction and no ATAC-seq peak. The remaining 74 enhancers (53%) were not predicted to have cell type-specific activity by either ATAC-seq or CREsted. Thus, considering DNA sequence along with ATAC-seq signal can help avoid testing inactive enhancers.

To illustrate the rescoring process, we show two enhancers that had scATAC-seq, CREsted and H3k27ac scores that were specific to oligodendrocytes (Oligo). First, an On-Target Oligo enhancer was predicted by the CREsted model to include two important candidate TF binding sites: one for the established Oligo TF SOX10 ^24,25^, and one for CREB5, known to play an important role in oligodendrocyte myelin synthesis and differentiation (**Fig. 4C**). Second, a No-Labeling enhancer was rescored as weakly On-Target for Oligos, and this was consistent with predictions from all modalities and a SOX10 binding site (**Fig. 4D**). To further understand sequence patterns that define cell type-specific On-Target enhancers, we calculated contribution scores for all On-Target enhancers in their corresponding cell types and identified frequently occurring motifs^26^ and clustered them across cell types ^18^ (**Supplemental Fig. 14**). For example, we found DLX-and LHX-like motifs in interneurons, MEF-like motifs and E-box motifs in neurons, SOX-like motifs in Oligo and Oligo precursors (OPC) and NFI-like motifs in astrocytes, Oligo, OPC, Vip and L6b neurons, consistent with previously reported TF motif enrichments in cortical cell types ^9^.

Using the rescored enhancers, we assessed the multi-label classification performance of enhancer activity by calculating precision and recall at different score thresholds. ATAC-seq modalities scored higher (average precision, AP = 0.51 using peak-scaled normalization, AP=0.52 using ArchR ReadInTSS normalization) than the CREsted sequence model (AP = 0.42) and H3k27ac (AP = 0.25) (**Fig. 4E**). A receiver operating characteristic (ROC) curve highlights the same trends for the different modalities (**Supplemental Fig. 15**). Interestingly, combining outputs from the CREsted and scATAC-seq models improves prediction (AP = 0.54), demonstrating that the modalities contain complementary information.

Next, we examined if the CREsted model captured sequence differences that were associated with the magnitude of *in vivo* activity of On-Target enhancers. On average, the model predicted higher scores for strong versus weak enhancers for 9 out of 14 cell types (**Fig. 4F**, **Supplemental Fig. 16**) in contrast to magnitude of the ATAC-seq signal (**Supplemental Fig. 17**). Additionally, we investigated how well models identified explainable negative enhancers. The CREsted model significantly (P < 0.05) outperformed scATAC approaches in avoiding false positives (**Fig. 4G**) and false negatives (**Supplemental Fig. 18**). Overall, these analyses underscore the benefits of using sequence models to improve non-functional enhancer prediction, prioritize strong enhancers, and lay the groundwork for understanding cell type enhancer codes.

## Discussion

The BICCN Challenge provides a valuable benchmark for future computational methods aimed at predicting functional cell type-specific enhancers. With 677 enhancers validated on a standardized pipeline and scoring criteria, this represents the largest collection of its kind ^6^. Importantly, both data and code are publicly available, facilitating further research and method refinement. However, limitations exist, including use of bulk ATAC-seq datasets and selection of enhancers near cell type markers that may not represent the most specific peaks per cell type. In addition, the validation experiments used PHP.eB-pseudotyped AAV that infects most mouse cortical cell types, although with biases^6,27^, and caution should be exercised in using these data to refine prediction models. Furthermore, the epifluorescence scoring, while valuable, is an imperfect estimate of target specificity based on cell morphology clues and spatial distributions of labeled cells. Finally, SSv4 data was collected from mouse primary visual cortex (V1), and validation results may not always align with model predictions based on mouse M1 multi-omic data, although cell subclasses share similar molecular profiles across the mouse neocortex ^7,28^.

The validation data from this challenge reveal which features are critical for predicting enhancer function, and the highest-performing methods primarily leveraged the specificity of ATAC-seq peaks in the mouse data set. Interestingly, the top submission combined RNA-seq with ATAC-seq to predict cell type-specific TF-enhancer-gene triplets. Additionally, incorporating HiC data was valuable for predicting enhancers in specific neuron types, such as L6b. H3K27ac and ABC scores tended to be higher in On-Target enhancers, although with lower recall and precision compared to cell-type specific ATAC-seq signals. Moreover, biological priors played a crucial role in the success of these models. For instance, including moderately sized ATAC-seq peaks was critical to predict candidate enhancers since the largest peaks often represent promoters.

Despite initial expectations that deep learning and DNA sequence models would outperform, the results showed that simpler ranking of ATAC-seq peaks by their cell type specific signal performed similarly. However, the top model did not consistently excel across all cell types and enhancer categories. The distinct methodologies used by the top three teams indicate that there are opportunities to refine enhancer prioritization. For example, the CREsted sequence model avoided Mixed-Target or Off-Target enhancers and predicted a strong oligodendrocyte enhancer by finding a coactivating TF motif AP-1 near a motif for SOX10, an oligodendrocyte marker ^24,25^. We further showed that combining cell type-specific predictions from the sequence model together with cell type-specific peaks from scATAC-seq data provides the best predictive factor of enhancer function and specificity (**Fig. 4E**). Future challenge submissions that incorporate more diverse molecular profiling of cortical cells will be critical in evaluating the importance of open chromatin data for enhancer prediction.

Looking forward, a major goal of the field is to develop a comprehensive toolkit to target cell types across the brain. Improving data coverage and quality is crucial, as many On-Target enhancers were missed due to low ATAC-seq signal. Fortunately, on-going work supported by the NIH BRAIN Initiative is focused on whole-brain atlasing of cell types, including high read-depth single cell ATAC-seq, HiC, and histone marker profiling. Sequence models will enable the rational design of enhancers tailored to cell types or groups of types, a strategy successfully applied in fruit fly models ^20^. Cross-species sequence models were under-represented in this challenge, but On-Target enhancers tend to have conserved open chromatin and sequence, and TF regulatory networks show conservation across primates and rodents ^1^. Additionally, sequence models trained on ATAC-seq data from multiple species enhance the prediction of chromatin accessibility^13^. Given these observations, we propose that decoding conserved enhancer codes presents a promising avenue for developing robust and precise tools to target cell types across many species.

It will be essential to expand validation experiments to include more brain regions and cell types and to quantify specificity and completeness of labeling using automated scoring of whole brain imaging and single cell sequencing. Community sharing of raw data will enable re-evaluation with future algorithms and comprehensive whole-brain analysis, while negative results will provide valuable insights into DNA repressor codes and enhancer function. Cross-species testing, including in non-human primates, will help assess the predictive power of these models across model organisms and will bolster confidence in translational applications for humans.

Integrating modeling and experimental approaches will be key to advancing enhancer tool development. Selecting enhancers that offer the most informative data for models will refine predictions and improve success rates. Testing enhancers with conflicting predictions from different modalities, such as ATAC-seq versus DNA sequence, could yield critical insights, even if they do not result in the most effective tools. Additionally, reinterpreting experiments with no labeling that have strong model support may uncover challenges in transducing some cell types such as has been reported for microglia ^29^. Ultimately, we need interpretable models that help to generate viral tools to target cell types and provide insights into cell type identity and genetic regulation in the context of human specializations and disease.

## Acknowledgements

This publication was coordinated through the NIH BRAIN Initiative Cell Census Network (BICCN) and Armamentarium for Precision Brain Cell Access (https://braininitiative.nih.gov/armamentarium). This work was funded by the Allen Institute for Brain Science and by NIH grants RF1MH121274 to B.T., 1UF1MH128339-01 to B.T., T.E.B., T.L.D., B.P.L. and J.T.T., R01MH113005 to R.L. and J.G., and RF1MH114126 and UG3MH120095 to E.S.L., J.T.T., and B.P.L. FWO PhD fellowship strategic basic research to N.K. (1SH6J24N). FWO PhD fellowship fundamental research to S.D.W. (1191323N). “Pioneer” and “Leading Goose” R&D Program of Zhejiang (2024SSYS0032) to K.Z. The authors thank the founder of the Allen Institute, Paul G. Allen, for his vision, encouragement and support.

## Author contributions

Viral tool testing: BPL, BT, DD, JKM, JTT, MH, TLD, YB Data analysis: BL, DA, DD, DM, ECE, EJA, FW, GH, GP, JDM, JG, JTT, KZ, MH, NH, NJJ, NK, NRZ, RL, SA, SDW, SS, TEB, VK, YB, YT Data interpretation: BL, BR, BT, DA, DD, DM, ECE, EJA, ESL, FW, GH, GP, JDM, JG, JM, JRE, JTT, KZ, MH, NH, NJJ, NK, NRZ, RL, SA, SDW, SS, TEB, VK, YB, YT Writing manuscript: BL, BT, DD, FW, JG, JTT, KZ, MH, NJJ, NK, NRZ, RL, SA, SS, TEB, YB, YT

## Data and materials availability

Cortical cell type AAV-based tools are described in Ben-Simon et al.^6^ and are available from Addgene (https://www.addgene.org/collections/brain-armamentarium).

## Methods

### Single nucleus molecular profiling

10x multiome ATAC + Gene Expression and methyl-3C-sequencing (snm3C-seq) experiments were carried out on the same tissue samples from human, macaque, marmoset, and mouse M1, as described ^13^.

### *In vivo* enhancer validation data

Detailed experimental methods are described in the companion paper ^6^ and other recent viral tools publications ^4,5,25^.

### Primary screen scoring

Each enhancer vector was screened and scored based on the labeling pattern it produced across the entire brain, with additional emphasis on cortical populations. First, each region of the brain where labeling of cell somata was observed was manually scored based on the labeling brightness and density, classifying each into either low or high. In addition, we created 11 categories of cell populations within the neocortex that could be visually distinguished one from the other. Whenever labeling was observed in one or more of these cortical populations, each population was individually evaluated based on its own brightness and density. Whereas brightness was classified based on whether the labeling was stronger or weaker than the common brightness observed across all experiments, density was evaluated based on the expected density of cells for each of the scored regions or populations, using the nuclear markers as reference. To determine target specificity, we aligned each target cell population with the labeled population which best matches its known anatomical location, distribution, and morphological characteristics. We determined an enhancer to be “On-Target” if the target population aligned with the labeled population, “Mixed-Target” if labeling was observed in populations other populations, in addition to the target one, “Off-Target” if labeling was observed exclusively in population/s other than the target population, and “No-Labeling” if no labeling was observed in the neocortex, regardless of whether labeling was observed in other brain regions.

### Cell type enhancer activity quantified by single cell RNA-seq

SSv4 data for cortical enhancers were generated from mouse primary visual cortex (V1/VISp) and mapped to the Allen Institute AIT2.1.1 VISp taxonomy as described in Ben-Simon et al. ^6^. The data were reused in this study to establish ground truth cell type specificity in combination with primary screening expression analysis.

### Benchmark metrics for evaluation of enhancer prioritization methods

We defined an interpretable benchmark metric based on the experimentally validated enhancer collection reported in Ben-Simon et al. ^6^. This benchmark metric is available to the community at: https://github.com/AllenInstitute/EnhancerBenchmark. The benchmark metric is a composite of an epifluorescence imaging score and SSv4 quantifications score per enhancer that captures different properties of enhancer activity. Epifluorescence validation of an enhancer captures broad patterns such as cortical layer labeling and provides cell type-specificity when cellular morphologies are known. Additionally, the intensity of SYFP2 fluorescence provides a measure of enhancer strength. We scored each validated enhancer based on the cell type-specific rank and placed more weight on strong enhancers being enriched in the top ranks. However, the epifluorescence validation does not provide a quantitative measure of specificity for each cell type targeted by the enhancer virus. To achieve this, we used SSv4 to quantify the transcriptome of cells with high SYFP2 fluorescence and mapped these cells to a cortical taxonomy to determine accurate abundances of each cell type targeted by the enhancer.

The epifluorescence metric was designed to be minimized when teams arranged On-Target enhancers in the top ranks per-cell type and Mixed Target enhancers relatively lower down the ranks as these enhancers both target the intended cell type as well as additional unintended cell types. Negative enhancer categories including Off-Target and No-Labeling enhancers optimize the metric when placed at the bottom of the ranked lists per-celltype. For each validated enhancer *E_i_* ∈ *E_validated_* in the ranked list across cell types we multiply the predicted rank *R_E_i__* of the enhancer with an indicator variable *I_Category_* which encodes enhancer categories (On-Target, Mixed-Target, Off-Target or No-Labeling). Then the metric is weighted by an indicator variable *I_Strength_* encoding the strength of SYFP2 epifluorescence (Strong, Weak or None).

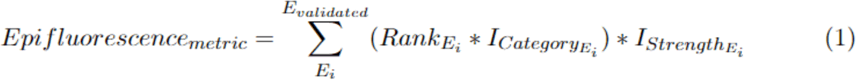

where:

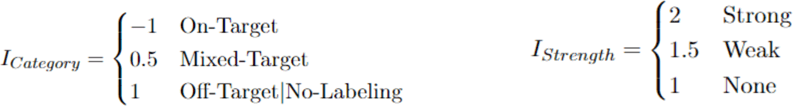

The SSv4 metric was designed to be minimized and replaces epifluorescence strength *I_Strength_* with a quantification of cell type-specificity computed the fraction of cells labeled as the intended target cell type 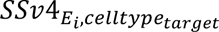 [0 − 1]. We then compute the SSv4 metric for each validated enhancer *E_i_* ∈ *E_validated_* by multiplying the predicted rank *R_E_i__* with the associated category *I_Category_* and the fraction of cells labeled as the targeted cell type 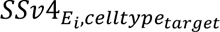.

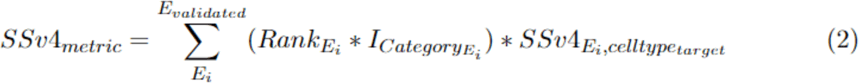

The composite benchmark score is then computed as the unweighted summation of Epi_metric and SSv4_metric.

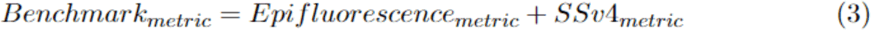

To define a normalized benchmark score ranging between 0 (worse) and 1 (better) we created an optimally ranked enhancer list per cell type and computed the associated benchmark metric. Then the benchmark metric score is divided by this optimal score and subtracted from 1 to achieve a normalized metric with the desired directionality.

To prevent overfitting during the challenge, we held back enhancers targeting excitatory cell types to ensure that no team could gain an advantage by learning patterns in the hidden enhancer validation data over the course of multiple challenge submissions.

### Statistical quantification of benchmark metric per-team

To assess the significance of the ranking of the given methods we set up a statistical test in which we defined:

- **Null Hypothesis:** The rank of the team is unstable and occurs uniformly across the possible ranks.
- **Alternative Hypothesis:** The rank of the team is stable (e.g., significantly clustered around its observed rank).

In which significantly clustered is defined as +/- 2 ranks of the teams rank from the full benchmark score. P-value on rank stability per-team were then empirically computed by randomly subsampling to 90% the validated enhancer database, 10,000 times.

### Recovery curves of functional enhancers per method

To assess the recovery of validated cell type specific enhancers based on the cell type specific rankings of each submission a recovery approach was used. In brief, the set of ground truth cell type specific enhancers was defined as the genomic regions (candidate enhancers) for which the specificity was classified as “On-Target”. Mouse genomic coordinates were used for candidate enhancers originating from the mouse genome, and coordinates lifted over to the mouse genome were used for candidate enhancers originating from the human genome. The genomic regions in each cell type specific ranking were intersected with the ground truth cell type specific enhancers using pyranges identifying hits along the ranking (genomic regions in the ranking overlapping with multiple ground truth enhancers were only counted once). Recovery curves were drawn for each submission by calculating the cumulative sum of the union of hits along all cell type specific rankings per cell type. In cases where the ranking was shorter than 10,000 elements, the ranking was padded with non-hits up to a length of 10,000 elements. Normalized enrichment scores (NES) per cell type specific ranking per submission were calculated as the area under the curve of the recovery curve (AUC) up to the 1,000th element and dividing this by the AUC up to the 1,000th element of the average recovery curve of 100 random rankings.

### Manual annotation of validated enhancers

We manually inspected a subset of the validated regions by classifying them into five categories: explainable positives, unexplainable positives, explainable negatives, unexplainable negatives, and undetermined. Explainable positives are defined as functional (both ‘strong’ and ‘weak’, and both ‘On-Target’ and ‘Mixed-Target’) enhancers that have either a strong accessibility peak in the scATAC-seq data, a strong prediction from the CREsted model, or both, in their intended target cell type(s). Unexplainable positives are functional enhancers that do not have a clear peak and/or prediction in their target cell types, but still show activity for those cell types, or they do have a strong peak and/or prediction in their targets, but show activity in other cell types. For validated enhancer candidates that did not show functionality (negatives), we identify explainable negatives as regions that either do not have a specific peak, a specific prediction, or both, in the target cell type. Unexplainable negatives have either a peak, prediction, or both in their target cell types, as well as the presence of positive contribution scores in one or more motifs obtained from the CREsted model. Undetermined regions do not contain enough decisive information to classify them into one of the other four categories, often because of their target cell types not being included in the original dataset.

### Precision-recall curves of different modalities

To compare the different modalities directly on the 677 validated enhancers, we scored each enhancer per modality by taking per cell type the score over the sum of all scores. We generated a target binary matrix based on the On- and Mixed-Target and No-Labeling enhancers, combined with the SSv4 results, to label the targeted cell types on enhancer functionality, and calculated per cell type the precision and recall over different prediction thresholds. We excluded off-target enhancers because of uncertainty for the actual targeted cell types.

For the scATAC-seq and H3k27ac data, we took the average counts over the exact region. The peak-scaled scATAC data was obtained by scaling the peak heights per cell type through the normalization factor obtained from the CREsted package. We then applied the specificity metric to the resulting target vectors per modality.

For the CREsted model the enhancer regions were put in the center of a 2114 bp background region, since the model takes in a fixed 2114 bp sequence. Then, we took the predictions scores per enhancer and applied our specificity metric to obtain final scores. The CREsted model is available at https://crested.readthedocs.io/en/latest/models/biccn.html.

For the combined CREsted and scATAC prediction scores, we averaged their specificity scores and recalculated the average precision and recall.

### Motif enrichment in On-Target enhancers

We calculated the most frequently occurring patterns per cell type in On-Target enhancers with *tfmodisco-lite*, based on contribution scores obtained from the CREsted model (Supplemental Fig. 14). We then matched patterns across cell types with the CREsted pattern matching function *crested.tl.modisco.process_patterns* (sim_threshold=4.25, trim_ic_threshold=0.1) We used the frequency of the patterns per sequence as *pattern_parameter* per cell type, and plotted the results using the *crested.pl.patterns.clustermap_with_pwm_logos* function with the importance threshold set to 0.12.

### Random forest modeling using biological priors and per enhancer genomics measures

To examine the importance of additional variables determining enhancer specificity, we modeled per enhancer metrics, including distance to the nearest transcription start site, mouse-human enhancer sequence conservation, and published ChIP seq data. ChIP seq data that overlapped enhancers were found using the GenomicRanges package in R, and all overlapping bins for each enhancer were summed to get values used for downstream analysis. Random forest models were constructed after removing enhancers that were ‘Mixed Target’ using scikit-learn version 1.3.0. The “% matching bps” measure was determined by calculating the percentage of the initial mouse enhancer length that is preserved upon examination of the best BLAST alignment with the human genome. For random forest model development, we used a 70/30 split for training and a held-out test set, respectively. Prior to testing on the held-out test set, models were constructed and validated using 10 fold cross-validation using GridSearchCV. The importance and statistical significance of variables is reported in Figures 3D and S10. Statistical significance was calculated using ANOVA, followed by a Tukey post hoc test.

### Stein Aerts team methods

#### ATAC-based rankings

We investigated the performance of rankings purely based on scATAC-seq data. We processed fragment files through pycisTopic ^17^:, a tool that aggregates counts in cells per cell type (subclass level) and CPM-normalizes them to account for the variability in cell numbers per type. For all consensus peaks, we obtained the mean accessibility per cell type from the pseudobulked cell type-specific accessibility tracks. We ranked regions based on the Gini index of their accessibility profile over all cell types to obtain the highest and most specific peaks per cell type. Using the Gini index only provides one value per region, so we assigned that value to the cell type with the highest peak value and gave a zero score to all the other cell types in that region. To further optimize this approach, we augmented the scATAC-seq data by merging the provided mouse dataset with publicly available mouse motor cortex datasets ^17,30^. We merged datasets by manually matching corresponding cell types, and weighted peak heights across datasets by using the number of cells per matched cell type.

#### Normalization of peak heights across cell types

Since peak heights across cell types are not always in the same scale after count normalization because of a potential difference in the total amount of accessible regions, we implemented an additional normalization method to create peak-scaled scATAC tracks across cell types. We used the CREsted peak normalization functionality with default parameters, a method that is aimed to alleviate this problem. This method takes thetop 1% (based on peak strength/height) of peaks per cell type, and only retains peaks which are generally accessible (Gini index < 0.25). This is done based on the assumption that strong generally accessible peaks should be in a similar range across all cell types. Based on those strong, generally accessible peaks, we compared the mean peak height per cell type and calculated scalars per cell type that ensure all mean peak heights become equal. These scalars are multiplied with the peak heights of their corresponding cell types to put peaks across different cell types in a more comparable range.

#### Sequence-based deep learning models

We trained two types of convolutional-based sequence-based accessibility prediction models from the CREsted package. The first is a peak regression model, a 9-layer convolutional neural network (CNN) model, trained on scATAC-seq data for chromatin accessibility analysis. It uses 2,114 bp DNA sequences as input and predicts the average accessibility signal per cell type on the center 1,000 bp of that region, inspired by ChromBPNet^31^. As a loss function for these models, we chose to take the sum of the mean squared error (MSE) and the negative cosine similarity (‘CosineMSE Loss’ in CREsted). We first pretrained these models on all the given consensus peaks, and we further finetuned them on differentially accessible regions (DARs). DARs were determined through different methods, through regions which had a ratio higher than 2 between their highest and second highest peak scalar value, or by taking regions which had a Gini index higher than the mean plus one standard deviation of all regions in the peak set. The finetuning ensures that the models learn cell-type specific features, since most consensus peaks are not specifically accessible in a low number of cell types. A second approach is topic classification, which was the modeling method used in previously published models^9,32–34^. We analyzed the scATAC-seq data using pycisTopic ^17^ to obtain topics per region required for training such models. Transfer learning to DARs made it possible to obtain cell type-specific predictions, as was done in Hecker & Kempynck et al. 2024 ^9^.

We mostly focused on the mouse scATAC-seq data for training our model. For the regression models, we also trained a model on the human scATAC-seq data and did cross-species predictions to obtain the mouse rankings required for the challenge. For the topic models, we trained a model on all four species. We did topic modeling after integrating all four datasets, to obtain topics representing shared regulatory features. We finetuned that model on mouse DARs.

We used two methods of ranking all the consensus peaks for these models. The first one was using the Gini index on the prediction scores over the cell types per region. The second one was calculating per region, for each prediction per cell type, the product of the difference between a given cell type prediction and the highest prediction in any other cell type, and the ratio between them. This gave a specificity ranking per cell type, per region, which could be sorted to generate global rankings.

#### Pattern based enhancer scoring

We reasoned that regions with heterogeneous motif content were more likely to be functional enhancers. For this purpose, we generated SHapley Additive exPlanations (SHAP) values for the top 10,000 regions per cell type from the CRESted model, generating explanations for the prediction score of that cell type. Of these, the top 5,000 regions per cell type were used to identify recurring, important patterns using the tfmodisco-lite package ^35^. To rank regions based on motif diversity, we calculated the Shannon diversity index based on the pattern hits. This index was multiplied by a signal over noise metric that was defined, per region, as the number of seqlets (short stretches of DNA with a high importance score) having a pattern hit divided by all seqlets.

#### Scenic+ triplet scores

We ran SCENIC+ on the provided mouse multiome dataset (scRNA-seq + scATAC-seq) using default parameters^17^. Based on these results we generated a ranking of all transcription factor (TF)-region-gene triplets, which is the aggregated ranking (see below) of the TF- and region-to-gene importance scores and TF-to-region ranking based on the motif score of all motifs annotated to that TF. To make this ranking cell type-specific we combined it with the gini-based cell type-specific ATAC ranking.

#### Aggregate rankings of multiple methods

To combine multiple rankings of different methods, we used OrderStatistics^36^. In brief, rankings were generated based on the score of each method, ties were broken by assigning incremental rankings to tied regions based on the order in which they happen to occur. Next, rank-ratios were calculated as the ranking divided by the number of regions in each ranking and combined in a single ranking using the formula described in and implemented in the SCENIC+ package ^17^.

### Jesse Gillis team method

We employed a multi-modal approach to predict cell-type-specific enhancers in the mouse brain, utilizing three distinct data types: single-cell RNA sequencing (scRNA-seq), single-cell Assay for Transposase-Accessible Chromatin sequencing (scATAC-seq), and meta-Hi-C. For scATAC-seq data, we obtained binarized MACS2 peak scores from three publicly available sources ^7,13,30^. These scores were then z-scored at each genomic bin to capture cell-type specificity accurately. To enhance the power of scATAC-seq data further, we aggregated the binarized peaks from the three scATAC-seq sources to create a robust scATAC-seq matrix. Again, this matrix was z-scored at each bin to enhance cell-type specificity.

In the case of meta-Hi-C, we used our in-house method to obtain cell-type-specific contact vectors by combining meta-Hi-C data with scRNA-seq marker genes. Previously, we aggregated hundreds of Hi-C contact matrices for mouse Hi-C to generate meta-Hi-C maps available at https://labshare.cshl.edu/shares/gillislab/resource/HiC/^37^. Similarly, we utilized scRNA-seq data to obtain reliable markers for each brain cell type in mice^38^. The cell-type-specific profile was derived by computing the mean of the top markers for each cell type and smoothing the Hi-C contact vector using various marker subsets. The resulting meta-Hi-C matrix was z-scored at each bin to enhance cell-type specificity. Only intra-chromosomal contact matrices were used for this analysis. The Hi-C vectors are slightly updated from the time of submission to the challenge, however this does not alter the validation results significantly.

To integrate the meta-ATAC-seq and meta-Hi-C data, we multiplied the ATAC-seq peak scores with the Hi-C contact scores for each cell type, resulting in the metaATAC X metaHi-C matrix. This matrix was also z-scored at each bin to capture cell-type-specific enhancers accurately.

### PeakRankR method

PeakRankR is an R function (https://github.com/AllenInstitute/PeakRankR) designed for prioritizing and ranking cell type-specific peaks based on chromatin accessibility data. It employs a linear model approach to calculate the combined effects of positive features associated with a given peak in a cell type, which are then used to rank the peaks.

The features computed by PeakRankR include:

1. Specificity: Calculated using an R version of ‘multiBigWigSummary’ from deeptools (https://deeptools.readthedocs.io/en/develop/).
2. Sensitivity: Determined by counting the number of cell types that have a peak at the genomic coordinates of the enhancer.
3. Magnitude: Estimated using MACS2 to assess the ATAC-Seq signal at the genomic coordinates of the enhancer for the cell type of interest.

These three features are combined to generate a score for each peak as a weighted linear sum of the features:

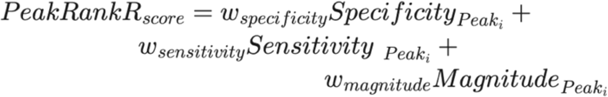

where *w* stands for the weight of each feature. By default, all weights are set to 1, indicating equal importance for each feature.

Function: Peak_RankeR (tsv_file_df, group_by_column_name, background_group, bw_table, rank_sum, weights)

- tsv_file_df: A tab-separated data frame of enhancer coordinates and peak groups.
- group_by_column_name: The column name in ‘tsv_file_df’ containing the groups of the enhancers.
- background_group: The group against which the enhancer should be prioritized for the group of interest.
- bw_table: A two-column table with the group BigWig file paths and group names.
- rank_sum: If TRUE, the sum of the feature scores is included in the output file.
- weights: Coefficients for the features, representing the influence of each predictor on the peak rank.

The function returns an object that includes the input peak coordinates, peak rank, and their corresponding scores.

- peakRankR_rank: Prioritized rank assigned to each peak
- rank_sum: Aggregate sum of scores for the various peak features

We enhanced PeakRankR to include additional features that characterize the shape of a peak: (1) Skew, which assesses the asymmetry of read pileups in an enhancer, with higher rankings given to those closer to symmetry (skewness near 0); (2) Kurtosis, which evaluates the dispersion of reads between the center and tails of an enhancer, with a higher rank for greater dispersion; (3) Modality, indicating the number of peaks within an enhancer, with unimodal peaks ranked higher. These features were calculated using functions from the ‘modes’ package in R (https://www.rdocumentation.org/packages/modes/versions/0.7.0).

Many peak ranking methods demand significant computational resources, and the chosen peaks may lack robustness. Therefore, a need exists for a straightforward and efficient function to rank peaks that can be adapted for specific cell types. PeakRankR addresses this by identifying the optimal features for a given peak set in a cell type and calculating a score based on specificity and sensitivity.

### Yoshiaki Tanaka team method

cisMultiDeep is the repository of R and Python scripts and command lines that identify functional CREs from single-cell multi-omics profiles. Given high conservation of the cell type-specific genes, we first obtained an orthologous gene list from Biomart (https://ensembl.org/info/data/biomart/) ^39^. On the other hand, the peak conservation was defined by UCSC LiftOver function in KentUtils and Bedtools (v2.30.0) ^40^. Subsequently, the cell type specificity in each gene and peak was estimated in RNA, mCG, mCH, and ATAC profiles by Wilcoxon rank-sum test. Here, the cell type specificity was calculated only in orthologous genes, whereas both conserved and non-conserved peaks were used for the assessment of the cell type specificity. In each cell type, the top 1,000 genes and 10,000 peaks were selected for subsequent deep learning analyses.

To ask if the selected genes/peaks are sufficient to define the cell types, we employed automatically-tuned deep neural network that was designed by Tensorflow python library (v2.9.0) with Keras Tuner API (v1.1.2) ^41,42^. Briefly, at first, dimensionality of the input data (RNA, mCG, mCH, and ATAC profiles) was reduced into 200 by principal component analysis (PCA). Then, the sequential neural network model was built with seven tunable hyperparameters: i) the number of layers (2 to 10 with increment of 1), ii) the number of nodes in hidden layers (50 to 500 with increment of 50), iii) dropout rates (0 to 0.5 with increment of 0.1), iv) activation functions (e.g. sigmoid), v) optimizers (e.g. Adam), vi) learning rate (e.g. 1e-1, 1e-2, 1e-3, 1e-4, 1e-5), and vii) loss functions (e.g. mean squared error loss function). Once the neural network model was optimized, mean absolute SHAP value, which represents the impact of each gene or peak on the cell type determination, was calculated by DeepSHAP that is a technique that can handle the complex and non-linear interactions across features and is optimized to calculate SHAP value for deep neural network ^43^.

Chromatin looping enables distal CREs to contact their target genes. Here, we hypothesized that the cell type-specific CREs are physically contacted with various cell type-specific genes. Thus, we ranked the peaks by the sum of the mean absolute SHAP values of the contacted genes by Hi-C loop. If the peak is conserved, we also added up the sum of the mean absolute SHAP values in other species.

### Kai Zhang team method

Using the genome annotation downloaded from Gencode, for each gene we extracted the 196,608 bp DNA sequences centered around its TSS. We then applied the Enformer model ^19^ to these sequences to derive sequence embeddings, followed by attention pooling to further reduce the dimensionality. Consequently, the processed sequences were represented as 896 vectors, each with 48 dimensions, corresponding to 128-bp segments of the initial sequence. Additionally, we quantified the ATAC-seq fragments overlapping these segments, adjusting for reads per kilobase million (RPKM).

To identify candidate enhancers, we developed a machine learning framework that integrates ATAC-seq signals and the aforementioned sequence embeddings to predict gene expression profiles. We first transformed normalized counts of ATAC-seq fragments into four-dimensional vectors using convolutional and self-attention layers. These ATAC embeddings were then merged with sequence embeddings and fed into a sequence of linear layers aimed at predicting levels of gene expression.

To train the model, we partitioned the data into training, validation and testing sets using an 80:10:10 split. We employed the Adam optimizer with a learning rate of 0.0001 and trained the model over 10 epochs, after which the model’s performance plateaued.

To determine the enhancer activity score for a specific region, we modified its ATAC signal by setting its count value to zero. The trained model was then used to make two predictions: one for the original (unmasked) input and another for the modified (masked) input. The enhancer activity score was calculated as the difference in the model’s predicted gene expression between these two inputs, serving as an indicator of the ATAC signal’s impact on the expression of adjacent genes. For each cell type, the top 10,000 highest scored peaks were chosen for submission.

#### ArchR team method

ArchR was run with default parameters following the tutorial: https://www.archrproject.com/bookdown. Briefly, we used ArchR to: (1) create pseudobulk replicates in which careful selection of background cells occurs; (2) assign of cell type labels to the ATAC-seq nuclei based on the RNA-seq component of the 10x Multiome; (3) call peaks for each cell type group; (4) use the getMarkerFeatures function to identify cell type specific peaks and associated summary stats; (5) organize 10,000 putative cell type-specific peaks for each cell type by sorting by log2FC > 2 and FDR < 0.05.

## References

1. Bakken, T.E., Jorstad, N.L., Hu, Q., Lake, B.B., Tian, W., Kalmbach, B.E., Crow, M., Hodge, R.D., Krienen, F.M., Sorensen, S.A., et al. (2021). Comparative cellular analysis of motor cortex in human, marmoset and mouse. Nature 598, 111–119. 10.1038/s41586-021-03465-8.

2. Yao, Z., Liu, H., Xie, F., Fischer, S., Adkins, R.S., Aldridge, A.I., Ament, S.A., Bartlett, A., Behrens, M.M., Van den Berge, K., et al. (2021). A transcriptomic and epigenomic cell atlas of the mouse primary motor cortex. Nature 598, 103–110. 10.1038/s41586-021-03500-8.

3. McColgan, P., Joubert, J., Tabrizi, S.J., and Rees, G. (2020). The human motor cortex microcircuit: insights for neurodegenerative disease. Nat. Rev. Neurosci. 21, 401–415. 10.1038/s41583-020-0315-1.

4. Mich, J.K., Graybuck, L.T., Hess, E.E., Mahoney, J.T., Kojima, Y., Ding, Y., Somasundaram, S., Miller, J.A., Kalmbach, B.E., Radaelli, C., et al. (2021). Functional enhancer elements drive subclass-selective expression from mouse to primate neocortex. Cell Rep. 34, 108754. 10.1016/j.celrep.2021.108754.

5. Graybuck, L.T., Daigle, T.L., Sedeño-Cortés, A.E., Walker, M., Kalmbach, B., Lenz, G.H., Morin, E., Nguyen, T.N., Garren, E., Bendrick, J.L., et al. (2021). Enhancer viruses for combinatorial cell-subclass-specific labeling. Neuron 109, 1449–1464.e13. 10.1016/j.neuron.2021.03.011.

6. Ben-Simon, Y., Hooper, M., Narayan, S., Daigle, T., Dwivedi, D., Way, S.W., Oster, A., Mich, J.K., Taormina, M.J., Martinez, R.A., et al. (2024). A suite of enhancer AAVs and transgenic mouse lines for genetic access to cortical cell types. bioRxiv, 2024.06.10.597244. 10.1101/2024.06.10.597244.

7. Zu, S., Li, Y.E., Wang, K., Armand, E.J., Mamde, S., Amaral, M.L., Wang, Y., Chu, A., Xie, Y., Miller, M., et al. (2023). Single-cell analysis of chromatin accessibility in the adult mouse brain. Nature 624, 378–389. 10.1038/s41586-023-06824-9.

8. Liu, H., Zeng, Q., Zhou, J., Bartlett, A., Wang, B.-A., Berube, P., Tian, W., Kenworthy, M., Altshul, J., Nery, J.R., et al. (2023). Single-cell DNA methylome and 3D multi-omic atlas of the adult mouse brain. Nature 624, 366–377. 10.1038/s41586-023-06805-y.

9. Hecker, N., Kempynck, N., Mauduit, D., Abaffyová, D., Vandepoel, R., Dieltiens, S., Sarropoulos, I., González-Blas, C.B., Leysen, E., Moors, R., et al. (2024). Enhancer-driven cell type comparison reveals similarities between the mammalian and bird pallium. bioRxiv, 2024.04.17.589795. 10.1101/2024.04.17.589795.

10. Yao, Z., van Velthoven, C.T.J., Kunst, M., Zhang, M., McMillen, D., Lee, C., Jung, W., Goldy, J., Abdelhak, A., Aitken, M., et al. (2023). A high-resolution transcriptomic and spatial atlas of cell types in the whole mouse brain. Nature 624, 317–332. 10.1038/s41586-023-06812-z.

11. Siletti, K., Hodge, R., Mossi Albiach, A., Lee, K.W., Ding, S.-L., Hu, L., Lönnerberg, P., Bakken, T., Casper, T., Clark, M., et al. (2023). Transcriptomic diversity of cell types across the adult human brain. Science 382, eadd7046. 10.1126/science.add7046.

12. Choobdar, S., Ahsen, M.E., Crawford, J., Tomasoni, M., Fang, T., Lamparter, D., Lin, J., Hescott, B., Hu, X., Mercer, J., et al. (2019). Assessment of network module identification across complex diseases. Nat. Methods 16, 843–852. 10.1038/s41592-019-0509-5.

13. Zemke, N.R., Armand, E.J., Wang, W., Lee, S., Zhou, J., Li, Y.E., Liu, H., Tian, W., Nery, J.R., Castanon, R.G., et al. (2023). Conserved and divergent gene regulatory programs of the mammalian neocortex. Nature 624, 390–402. 10.1038/s41586-023-06819-6.

14. Kaplow, I.M., Schäffer, D.E., Wirthlin, M.E., Lawler, A.J., Brown, A.R., Kleyman, M., and Pfenning, A.R. (2022). Inferring mammalian tissue-specific regulatory conservation by predicting tissue-specific differences in open chromatin. BMC Genomics 23, 291. 10.1186/s12864-022-08450-7.

15. EnhancerBenchmark (Github).

16. Granja, J.M., Corces, M.R., Pierce, S.E., Bagdatli, S.T., Choudhry, H., Chang, H.Y., and Greenleaf, W.J. (2021). ArchR is a scalable software package for integrative single-cell chromatin accessibility analysis. Nat. Genet. 53, 403–411. 10.1038/s41588-021-00790-6.

17. Bravo González-Blas, C., De Winter, S., Hulselmans, G., Hecker, N., Matetovici, I., Christiaens, V., Poovathingal, S., Wouters, J., Aibar, S., and Aerts, S. (2023). SCENIC+: single-cell multiomic inference of enhancers and gene regulatory networks. Nat. Methods 20, 1355–1367. 10.1038/s41592-023-01938-4.

18. Kempynck, N., Mahieu, L., Ekşi, E., Can, K., Vasilis, B., Cas De Winter, S., Hulselmans, G., and Aerts, S. (2024). CREsted: Cis Regulatory Element Sequence Training, Explanation, and Design. 10.5281/zenodo.13320756.

19. Avsec, Ž., Agarwal, V., Visentin, D., Ledsam, J.R., Grabska-Barwinska, A., Taylor, K.R., Assael, Y., Jumper, J., Kohli, P., and Kelley, D.R. (2021). Effective gene expression prediction from sequence by integrating long-range interactions. Nat. Methods 18, 1196–1203. 10.1038/s41592-021-01252-x.

20. Taskiran, I.I., Spanier, K.I., Dickmänken, H., Kempynck, N., Pančíková, A., Ekşi, E.C., Hulselmans, G., Ismail, J.N., Theunis, K., Vandepoel, R., et al. (2024). Cell-type-directed design of synthetic enhancers. Nature 626, 212–220. 10.1038/s41586-023-06936-2.

21. Creyghton, M.P., Cheng, A.W., Welstead, G.G., Kooistra, T., Carey, B.W., Steine, E.J., Hanna, J., Lodato, M.A., Frampton, G.M., Sharp, P.A., et al. (2010). Histone H3K27ac separates active from poised enhancers and predicts developmental state. Proceedings of the National Academy of Sciences 107, 21931–21936. 10.1073/pnas.1016071107.

22. Xie, Y., Zhu, C., Wang, Z., Tastemel, M., Chang, L., Li, Y.E., and Ren, B. (2023). Droplet-based single-cell joint profiling of histone modifications and transcriptomes. Nat. Struct. Mol. Biol., 1–6. 10.1038/s41594-023-01060-1.

23. Fulco, C.P., Nasser, J., Jones, T.R., Munson, G., Bergman, D.T., Subramanian, V., Grossman, S.R., Anyoha, R., Doughty, B.R., Patwardhan, T.A., et al. (2019). Activity-by-contact model of enhancer– promoter regulation from thousands of CRISPR perturbations. Nat. Genet. 51, 1664–1669. 10.1038/s41588-019-0538-0.

24. Stolt, C.C., Rehberg, S., Ader, M., Lommes, P., Riethmacher, D., Schachner, M., Bartsch, U., and Wegner, M. (2002). Terminal differentiation of myelin-forming oligodendrocytes depends on the transcription factor Sox10. Genes Dev. 16, 165–170. 10.1101/gad.215802.

25. Mich, J.K., Sunil, S., Johansen, N., Martinez, R.A., Leytze, M., Gore, B.B., Mahoney, J.T., Ben-Simon, Y., Bishaw, Y., Brouner, K., et al. (2023). Enhancer-AAVs allow genetic access to oligodendrocytes and diverse populations of astrocytes across species. bioRxivorg. 10.1101/2023.09.20.558718.

26. Shrikumar, A., Tian, K., Avsec, Ž., Shcherbina, A., Banerjee, A., Sharmin, M., Nair, S., and Kundaje, A. (2018). Technical note on Transcription Factor Motif Discovery from Importance Scores (TF-MoDISco) version 0.5.6.5. arXiv [cs.LG].

27. Shi, H., He, Y., Zhou, Y., Huang, J., Maher, K., Wang, B., Tang, Z., Luo, S., Tan, P., Wu, M., et al. (2023). Spatial atlas of the mouse central nervous system at molecular resolution. Nature 622, 552–561. 10.1038/s41586-023-06569-5.

28. Yao, Z., van Velthoven, C.T.J., Nguyen, T.N., Goldy, J., Sedeno-Cortes, A.E., Baftizadeh, F., Bertagnolli, D., Casper, T., Chiang, M., Crichton, K., et al. (2021). A taxonomy of transcriptomic cell types across the isocortex and hippocampal formation. Cell. 10.1016/j.cell.2021.04.021.

29. Okada, Y., Hosoi, N., Matsuzaki, Y., Fukai, Y., Hiraga, A., Nakai, J., Nitta, K., Shinohara, Y., Konno, A., and Hirai, H. (2022). Development of microglia-targeting adeno-associated viral vectors as tools to study microglial behavior in vivo. Commun. Biol. 5, 1224. 10.1038/s42003-022-04200-3.

30. Li, Y.E., Preissl, S., Hou, X., Zhang, Z., Zhang, K., Qiu, Y., Poirion, O.B., Li, B., Chiou, J., Liu, H., et al. (2021). An atlas of gene regulatory elements in adult mouse cerebrum. Nature 598, 129–136. 10.1038/s41586-021-03604-1.

31. Creators Pampari, Anusri Shcherbina, Anna Nair, Surag Schreiber, Jacob Patel, Aman Wang, Austin Kundu, Soumya Shrikumar, Avanti Kundaje, Anshul Bias factorized, base-resolution deep learning models of chromatin accessibility reveal cis-regulatory sequence syntax, transcription factor footprints and regulatory variants 10.5281/zenodo.10396047.

32. Minnoye, L., Taskiran, I.I., Mauduit, D., Fazio, M., Van Aerschot, L., Hulselmans, G., Christiaens, V., Makhzami, S., Seltenhammer, M., Karras, P., et al. (2020). Cross-species analysis of enhancer logic using deep learning. Genome Res. 30, 1815–1834. 10.1101/gr.260844.120.

33. Bravo González-Blas, C., Matetovici, I., Hillen, H., Taskiran, I.I., Vandepoel, R., Christiaens, V., Sansores-García, L., Verboven, E., Hulselmans, G., Poovathingal, S., et al. (2024). Single-cell spatial multi-omics and deep learning dissect enhancer-driven gene regulatory networks in liver zonation. Nat. Cell Biol. 26, 153–167. 10.1038/s41556-023-01316-4.

34. Janssens, J., Aibar, S., Taskiran, I.I., Ismail, J.N., Gomez, A.E., Aughey, G., Spanier, K.I., De Rop, F.V., González-Blas, C.B., Dionne, M., et al. (2022). Decoding gene regulation in the fly brain. Nature 601, 630–636. 10.1038/s41586-021-04262-z.

35. Schreiber, J. tfmodisco-lite: A lite implementation of tfmodisco, a motif discovery algorithm for genomics experiments (Github).

36. Aerts, S., Lambrechts, D., Maity, S., Van Loo, P., Coessens, B., De Smet, F., Tranchevent, L.-C., De Moor, B., Marynen, P., Hassan, B., et al. (2006). Gene prioritization through genomic data fusion. Nat. Biotechnol. 24, 537–544. 10.1038/nbt1203.

37. Lohia, R., Fox, N., and Gillis, J. (2022). A global high-density chromatin interaction network reveals functional long-range and trans-chromosomal relationships. Genome Biol. 23, 238. 10.1186/s13059-022-02790-z.

38. Fischer, S., and Gillis, J. (2021). How many markers are needed to robustly determine a cell’s type? iScience 24, 103292. 10.1016/j.isci.2021.103292.

39. Kinsella, R.J., Kähäri, A., Haider, S., Zamora, J., Proctor, G., Spudich, G., Almeida-King, J., Staines, D., Derwent, P., Kerhornou, A., et al. (2011). Ensembl BioMarts: a hub for data retrieval across taxonomic space. Database 2011, bar030. 10.1093/database/bar030.

40. Quinlan, A.R., and Hall, I.M. (2010). BEDTools: a flexible suite of utilities for comparing genomic features. Bioinformatics 26, 841–842. 10.1093/bioinformatics/btq033.

41. Abadi, M., Barham, P., Chen, J., Chen, Z., Davis, A., Dean, J., Devin, M., Ghemawat, S., Irving, G., Isard, M., et al. (2016). TensorFlow: A system for large-scale machine learning. arXiv [cs.DC].

42. O’Malley, et al. (2019). Keras Tuner. https://github.com/keras-team/keras-tuner.

43. Lundberg, S., and Lee, S.-I. (2017). A Unified Approach to Interpreting Model Predictions. arXiv [cs.AI].

