## Supplemental Text for "Evaluating Methods for the Prediction of Cell Type-Specific Enhancers in the Mammalian Cortex"

[1]Allen Institute for Brain Science, Seattle, WA 98109.
[2]VIB Center for AI & Computational Biology, VIB-KU Leuven Center for Brain and Disease Research & KU Leuven Department of Human Genetics, Leuven, Belgium.
[3]Center for Epigenomics, Department of Cellular and Molecular Medicine, University of California, San Diego, La Jolla, CA 92093.
[4]Physiology Department and Donnelly Centre for Cellular and Biomolecular Research, University of Toronto, Toronto, Ontario, Canada.
[5]Maisonneuve-Rosemont Hospital Research Centre, University of Montreal, Montreal, Quebec, Canada.
[6]School of Life Sciences, Westlake University, Hangzhou, Zhejiang, China.
[7]Bioinformatics and Systems Biology Program, University of California, San Diego, La Jolla, CA 92093.
[8]Westlake Laboratory of Life Sciences and Biomedicine, Hangzhou, Zhejiang, China
[9]Department of Physiology and Biophysics, University of Washington, Seattle, WA 98195.
[10]Salk Institute for Biological Studies, La Jolla, CA 92037.
[11]These authors contributed equally
[12]Lead contact

### Stein Aerts lab results

For the blind challenge, we first preprocessed the raw sequencing files and re-analyzed the human and mouse scATAC-seq data sets using cisTopic (based on fragment counts, **Methods**). This validated the provided cell type annotation with respect to 2D dimensionality reduction (**Supplemental Text Figure 1A**). Next, we employed four different strategies to generate 4 types of rankings (each type may have one or more variants), each containing 19 ranked lists of the top 10,000 scoring intervals, for each of the given 19 cell types. The first strategy uses only cell-type specific chromatin accessibility to prioritize candidate enhancers per cell type by calculating per region the Gini index over the accessibility profile of all cell types. As a variant of this approach we took the pseudobulk profiles per cell type are derived from three datasets [13,17,29], instead of only the provided mouse dataset, followed by the application of a peak-scaling factor (**Methods**). These ATAC-only models resulted in very high performances according to the BICCN challenge scoring (see **Supplemental Fig. 1, Supplemental Table 2**): 0.4027 for the single dataset ATAC rankings, the best ranking in the entire challenge, and 0.4052 for the merged datasets.

The remaining three strategies all use sequence-based models. In the second strategy (**Supplemental Text Figure 1C**), we trained two types of convolutional neural network models to predict, from the sequence of the genomic interval as input, chromatin accessibility across cell types: CREsted peak regression and topic classification. We also trained analogous models on the human scATAC-seq clusters and used them to score and rank the mouse genomic regions, followed by an order-statistics integration of the mouse- and human-based rankings (**Methods**). Among the sequence-based models, the best performing model is a peak regression model (referred to in main text as Aerts CREsted), trained only on the mouse scATAC-seq data. The 10k rankings sorted on Gini score based on the model's predictions had a final score of 0.3556. In the third strategy, we combined the scATAC-seq and scRNA-seq data and inferred enhancer gene regulatory networks (eGRNs) using SCENIC+ [17]. This provides, as one of its outputs, triplet scores for TF-region-gene trios (**Supplemental Text Figure 1B**). This resulted in the best score in the challenge: 0.4086. In the fourth strategy, we generated SHAP-based explainability profiles for each candidate region, followed by seqlet clustering, and re-annotation of the regions using TFMoDisco-derived patterns [34] (**Supplemental Text Figure 1D**). Ranking was then performed using a heterogeneity score that measures the complexity of these patterns in each sequence (**Methods**). Combined with the scATAC-seq rankings, this resulted in the second highest score in the challenge: 0.4052 Finally, we also combined multiple strategies, using order-statistics, but overall these ensemble rankings did not outperform the ATAC-only or the CREsted approaches.

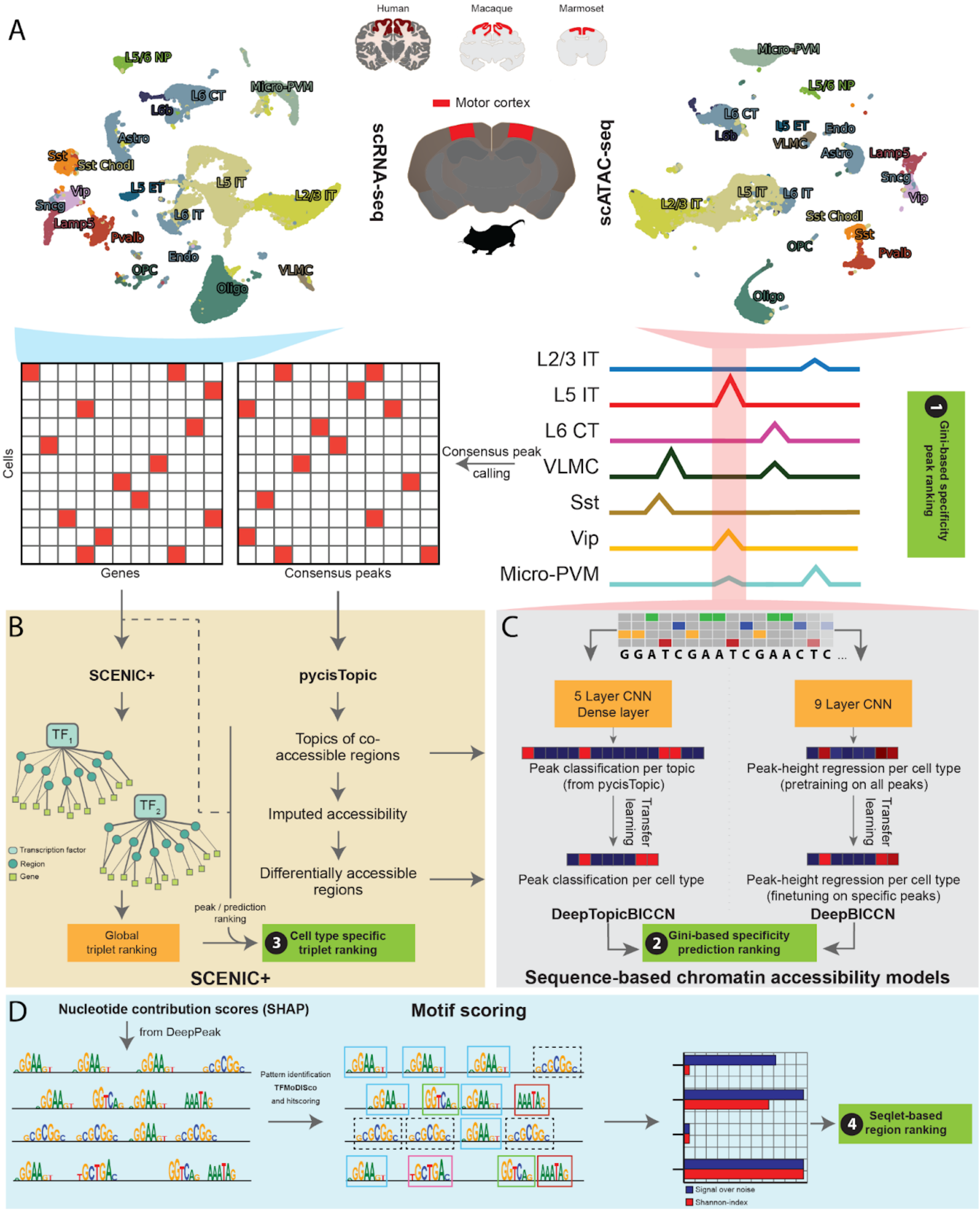


***Supplemental Text Figure 1: Overview of Aerts lab methods.***

**A.** Overview of challenge data from human and mouse using UMAP representation labeled by cell type. Schematic of data processing for subsequence methods. **B.** Schematic of the TF-region-gene trios SCENIC+ and topic assignment to peaks using pycisTopic. **C.** Schematic of DNA sequence models used to predict cell type-specific ATAC-Seq signal. **D.** Schematic of seqlet based methods which utilized motif scoring to determine enhancer specificity.

**Saroja Somasundaram lab results**

PeakRankR seeks to identify a minimal set of ATAC-seq features – namely, specificity, magnitude, sensitivity, and shape-related features like modality, skewness and kurtosis (**Methods**) – for each enhancer. Together, these features can be used to prioritize peaks simply and efficiently, like how experts select peaks using the UCSC genome browser. To assess the performance of PeakRankR, we investigated how well the method organized On-Target enhancers in the top predicted ranks and Mixed-Target, Off-Target and No-Labeling enhancers in lower ranks. Overall, PeakRankR performed comparable with or better than most other methods.

To evaluate the importance of each ATAC-seq feature in predicting the validated enhancers, we re-prioritized and evaluated peaks for the universe peak set using each PeakRankR feature separately. We find that specificity and magnitude using the bigWigSummary tool from UCSC optimize benchmark scores most efficiently, indicating that these aspects of peaks are the most important for prioritizing functional enhancers.

Additionally, PeakRankR is easily extendable in that additional peak features can be incorporated into the model's peak prioritization process. To determine whether peak shape impacts performance we additionally included modality, skewness and kurtosis (**Methods**) in a revised PeakRankR challenge submission. However, inclusion of these additional features did not improve upon the model leveraging specificity, magnitude and sensitivity. Identifying the most parsimonious set of features is needed for solving an optimization problem.

The lightweight and simple nature of PeakRankR enables rapid investigation of individual peak metrics and provides an efficient method to prioritize functional enhancers in the mammalian brain.

However, after the challenge concluded, we noted that PeakRankR utilized cell type calls and peaks filtered from ArchR as input. Therefore, PeakRankR was not provided with the complete set of ATAC-seq peaks, including some On-Target enhancers and cell types due to the filtered input. Inclusion of these peaks may have improved PeakRankR's score in the challenge. Scores for all PeakRankR submissions, both pre and post challenge, are included in the challenge GitHub repository.

**Jesse Gillis lab results**

We assessed the accuracy of our methods predictions using a set of experimentally validated enhancers from Ben-Simon et al. [6]. Our evaluation focused on determining the presence of cell type-specific enhancers based on evaluating accessibility and activity per cell type.

We evaluated our methods against the validation set using the area under the Receiver Operating Characteristic (AUROC) curve and Precision-Recall (PR) curve as performance metrics for this multi-label classification task (**Supplemental Text Figure 2**). Higher values of AUROC and PR area indicate stronger performance for each of our predictive models. We find that meta-ATAC-seq demonstrates marginally superior performance compared to leveraging individual ATAC-seq experiments, showcasing the predictive capability of robust ATAC-seq signals in identifying cell-type-specific enhancers. Interestingly, metaATAC combined with metaHi-C showed a marginal improvement over meta-ATAC-seq in PR curves. This suggests that incorporating ATAC-seq data along with contact density at enhancer regions is an important feature to enhancer prediction. Our finding aligns with observations from analogous activity by contact (ABC)[23] models for predicting gene expression. However, the ATAC-seq signal itself remains the most important feature when predicting cell-type specificity of enhancers. Additionally, we examined the AUC and AUPRC scores for each subclass and found that predicting enhancer specificity for non-neuron subclasses is relatively easier compared to neurons.

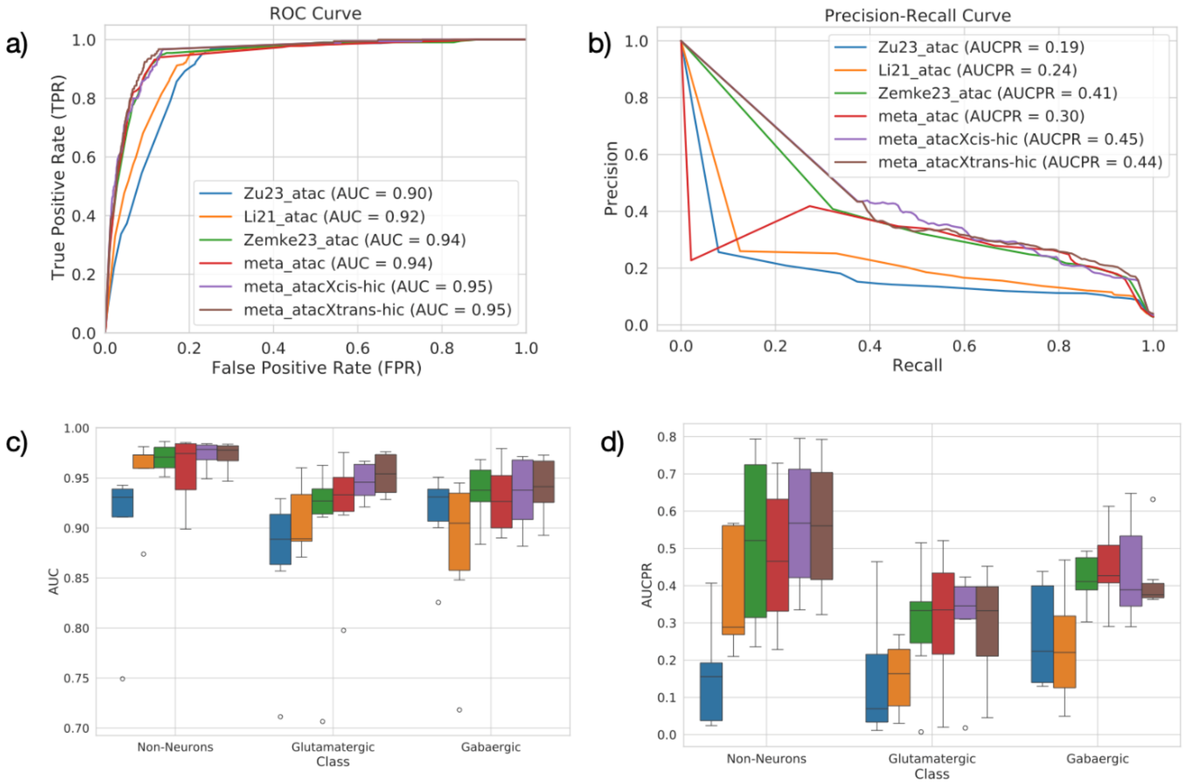


***Supplemental Text Figure 2: Assessment of Gillis lab methods against validated enhancers.***

**A.** Area under the receiver operator curve to assess performance of enhancer ranking methods using a reduced set of enhancers which were included in Gillis lab predictions. **B.** Performance of enhancer ranking methods using a reduced set of enhancers which were included in Gillis lab predictions. AP, average precision. **C.** Box plot of AUROC per method split by class (Glutamatergic, Gabaergic and Non-neurons). **D.** Box plot of AUCPR per method split by class (Glutamatergic, Gabaergic and Non-neurons).

**Yoshiaki Tanaka results**

The cell type is determined by expression of a specific set of genes that are governed by cis-regulatory elements (CREs), such as promoters and enhancers. CREs are characterized with open chromatin structure, in which transcription factors (TFs) are accessible. TF-bound CREs physically contact the target genes by forming chromatin loops, and transfer the regulatory information to the genes [43]. Importantly, the expression patterns of the cell type-specific genes and TFs are conserved across species [44,45], whereas cell type-specific CREs are more susceptible to evolutionary divergence [46,47] , although some highly regulatory CREs are conserved [48]. In addition, the cell type-specific gene expression also coincides with epigenetic modifications, such as DNA methylation (mCG and mCH). In particular, recent studies indicated that intragenic DNA methylation is negatively correlated with the gene expression [49,50]. These observations suggest that the cell type-specific transcriptional programs are associated with various determinants, including conservation, chromatin looping, and open chromatin, and the framework to integrate such multiple omics data is essential. Here, we introduce a pipeline, cisMultiDeep (https://github.com/ytanaka-bio/cisMultiDeep), that employs automatically-tuned deep learning with Shapley Additive exPlanation (SHAP) feature importance assessment in cross-species single-cell multi-omics data: RNA, ATAC, mCG, mCH, and Hi-C. First, this method identifies 'conserved' cell type-specific genes from RNA, mCG, and mCH profiles. Subsequently, 'conserved' and 'non-conserved' cell type-specific CREs are identified from ATAC profiles and linked with 'conserved' cell type-specific genes using deep learning. Finally, the contribution of each CRE to the cell type-specific gene expression is assessed by SHAP value and Hi-C contact map.

To evaluate our method we first intersected the validated enhancer collection with our top 10,000 CRE list. Amongst the 677 validated enhancers 27 peaks are unique to humans and do not have liftOver mouse genomic coordinates and were removed from subsequent assessment (**Supplemental Text Fig. 3A**). In addition, 131 peaks were also removed, since their coordinates were not overlapped with any mouse ATAC peaks from Zemke et al. 2023 [13]. in the remaining 519 validated enhancers, only 92 (17.8%) were detected in our top 10,000 CRE list. If we expanded the per-cell type ranked list to top 50,000 we detect 333 (64.1%).

To dissect the characteristics of CREs predicted by our model, we analyzed: (1) conservation, (2) magnitude, (3) distance from TSS, (4) specificity, (5) H3K27ac, and (6) ABC score[23]. We found that our model preferentially detected the conserved peaks ($p<3.68e-275$ by hypergeometric test) (**Supplemental Text Fig. 3B**) with high magnitude ($p<2.2e-16$ by T test) (**Supplemental Text Fig. 3C**). We note that the magnitude of the validated peaks was much lower than that of the predicted CREs ($p=7.95e-10$ by T test). Usually, ATAC peaks in proximal CREs to genes are likely to display higher magnitude than those in distant CREs. Thus, we also compared the distance of the predicted CREs from transcription start sites (TSSs), and found that our model preferentially identified proximal CREs, whereas many validated enhancers are distal CREs ($p<2.2e-16$ by T test) (**Supplemental Text Fig. 3D**). Furthermore, the rank of our top 10,000 CRE list was positively correlated with specificity and H3K27ac signal, but not ABC score (**Supplemental Text Fig. 3E**). Taken together, these assessments indicated that our model preferentially detected cell type-specific promoter elements, and requires refinement to capture distal CREs.

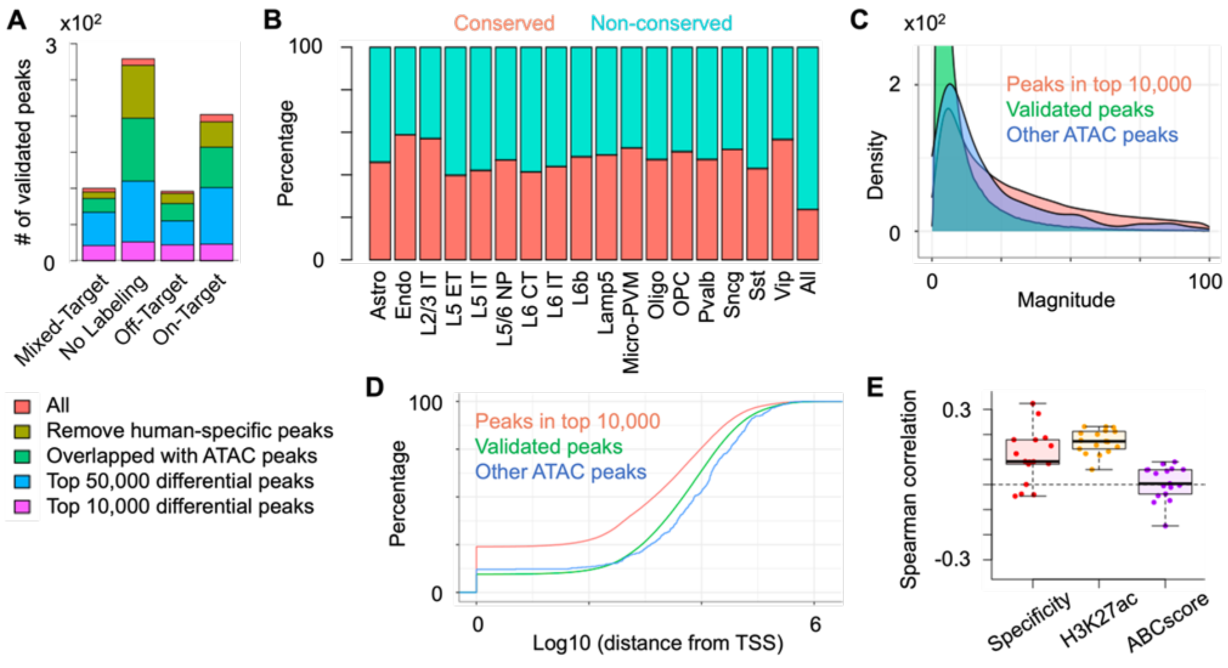


***Supplemental Text Figure 3. Assessment of cisMultiDeep model***.

**A.** Overlap of the validated peaks with the predicted CREs. **B.** Percentage of conserved ATAC peaks in top 10,000 CRE list. **C.** Density plot showing magnitude of our top 10,000 CREs, validated enhancers, and other ATAC peaks. **D.** Cumulative plot showing distance from TSS in our top 10,000 CREs, validated enhancers, and other ATAC peaks. **E.** Boxplot showing spearman correlation of specificity, H3K27ac and ABC score with the rank of our top 10,000 CRE list.

**Kai Zhang results**

We developed a machine learning algorithm leveraging the pre-trained Enformer model [19] to predict gene expression from DNA sequence and chromatin accessibility around TSS regions. Our model achieved great accuracy in predicting gene expression at varying abundance (PCC = 0.91, **Supplemental Text Fig. 4A**). To identify candidate enhancers, we performed in-silico perturbation of chromatin accessibility at each ATAC peak, and used the model to compute the changes of predicted gene expression before and after perturbation. These values were named enhancer scores. We used the experimentally validated cell-type-specific enhancers to assess the accuracy of our predictions (**Supplemental Text Fig. 4B**). We find that the average AUC in all cell types is 0.52. We selected the top 10,000 peaks with the highest enhancer scores as candidate enhancers in each of the 19 cell types. We found out that this set covered about 18% of the validated enhancers, and the coverage increased to 84% when we retained the top 100,000 peaks from each cell type (**Supplemental Text Fig. 4C**)

To further evaluate the correlation between enhancers and other information of peaks, we selected mouse enhancers with cell type specific ATAC-seq, identifying a total of414 enhancers. We calculated the AUC for these enhancers using ABC score (**Supplemental Text Fig. 4D**), H3K27ac signal (**Supplemental Text Fig. 4E**), and ATAC signal (**Supplemental Text Fig. 4F**) of peaks for different cell types, giving an average AUC of 0.82, 0.82, and 0.93, respectively. Of all the validation enhancers, 328 enhancers were located in the regions around genes' TSS.

---

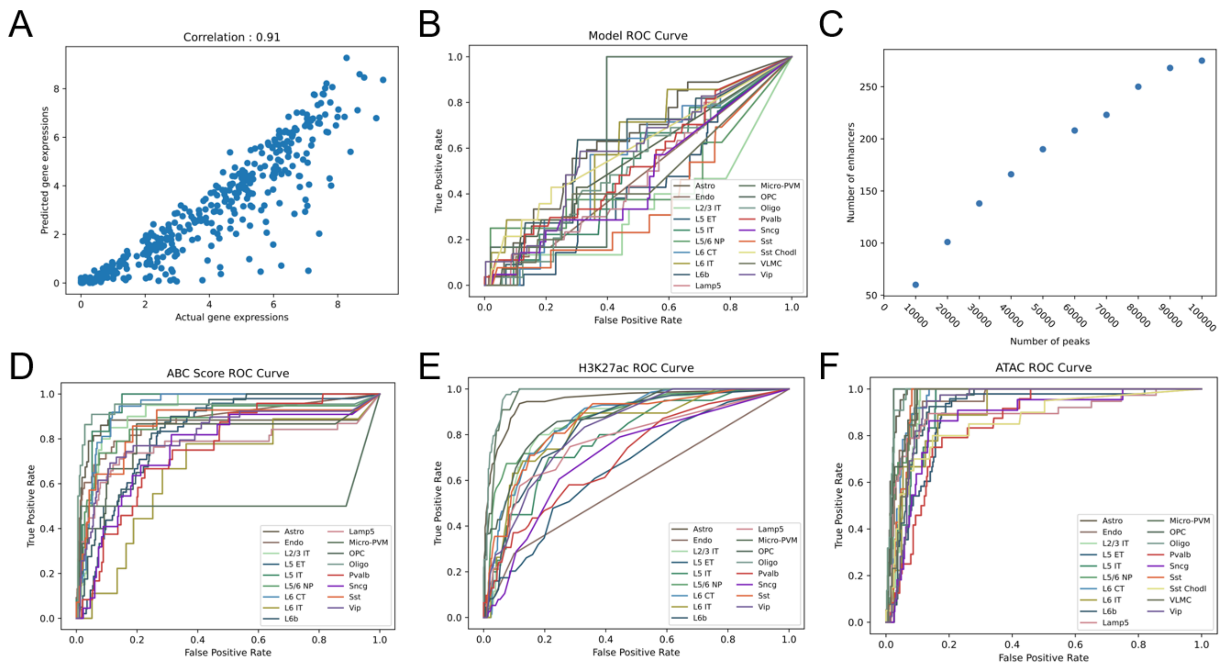


***Supplemental Text Figure 4. Assessment of an Enformer-based model.***

**A.** Scatter plot comparing predicted gene expression to actual gene expression. A total of 18,073 genes were downsampled to 400, ensuring an equal representation of genes across different expression ranges (0-2, 2-4, 4-6, >6). **B.** The ROC curve for 19 distinct cell types was generated using the enhancer score from our model. **C.** Scatter plot illustrating the number of experimentally validated enhancers captured by our predictions against the total number of enhancers predicted by our model. **D.** The ROC curve for 17 distinct cell types (excluding two cell types which were not accessible) was generated using the ABC scores of peaks. **E.** The ROC curve for 17 distinct cell types (excluding two cell types that were not accessible) was generated using the H3K27ac values of peaks. **F.** The ROC curve for 19 cell types was generated using RPKM ATAC signals of peaks.
