## Supplemental Figures and Table Legends for "Evaluating Methods for the Prediction of Cell Type-Specific Enhancers in the Mammalian Cortex"

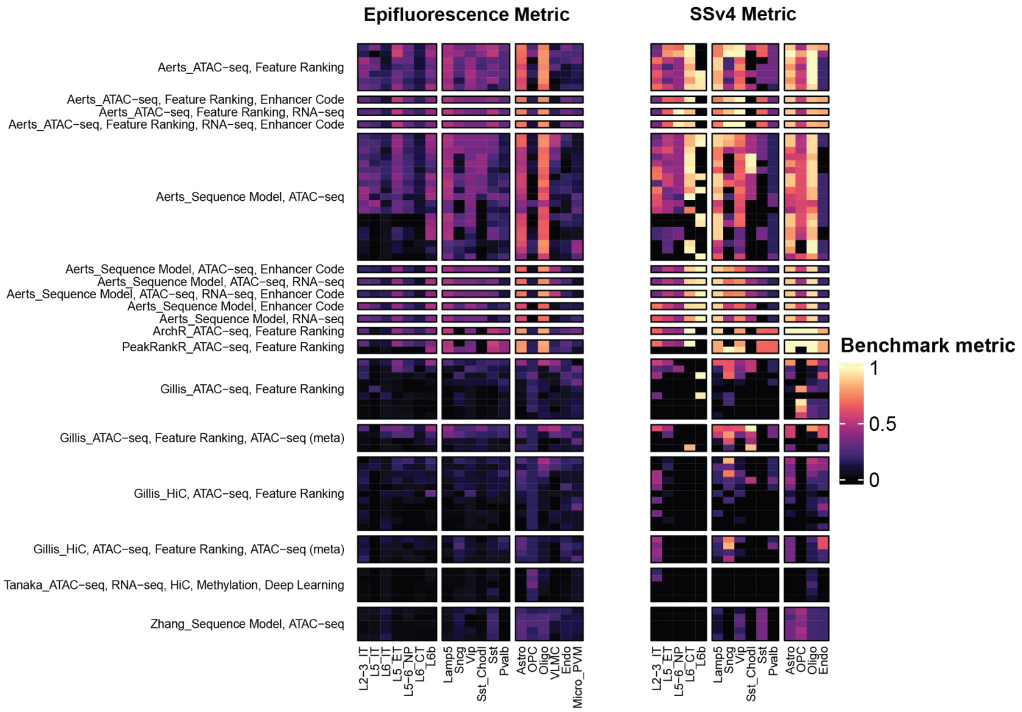


**Figure S1. Scoring of all team submissions based on two measures of *in vivo* enhancer activity.**
Submissions are grouped by team and approach. The heatmaps visualize the epifluorescence and SSv4 normalized benchmark metrics for each submission per subclass. Higher benchmark metric values represent agreement with the validated enhancer collection.

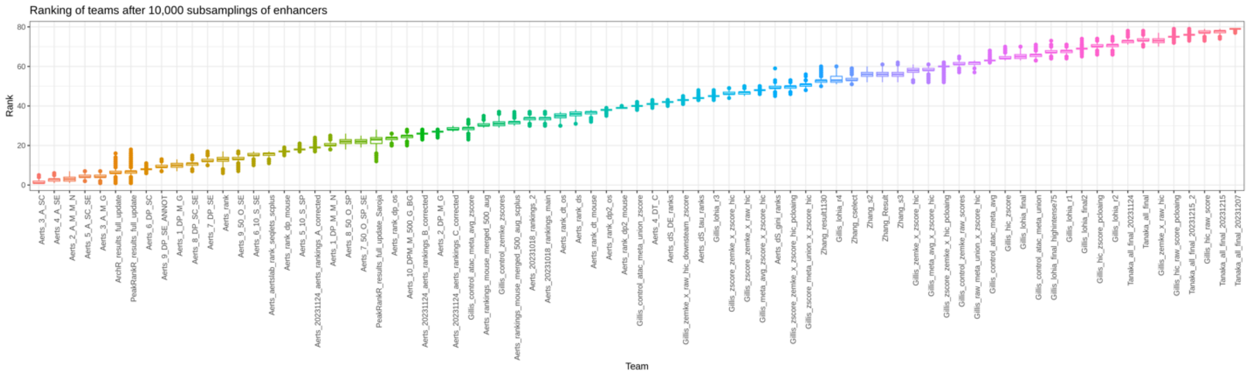


**Figure S2. Robustness of rank for each team submissions.**
Submissions are ordered by the final challenge benchmark metric in which teams aimed to achieve a low score. The box and whisker plots show the rank achieved by each submission across 10,000 random bootstraps of the validated enhancer database.

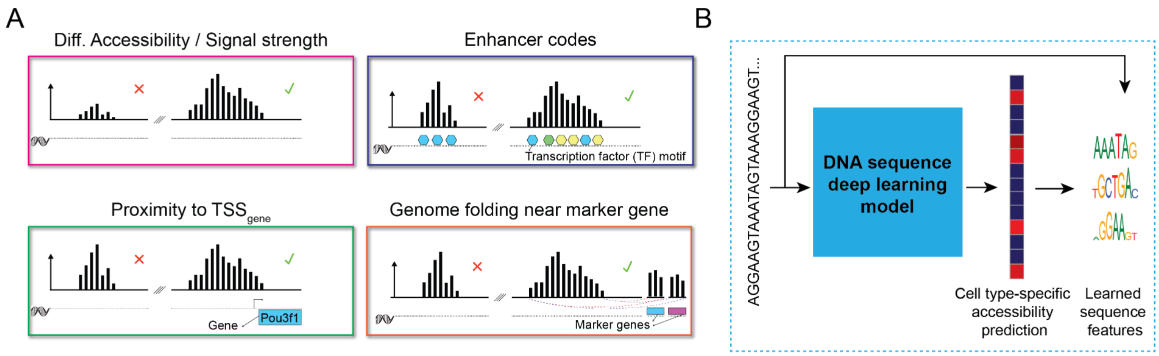


**Figure S3. Illustration of biological priors, methods and DNA sequence models**
Illustration of several genomic features and computational methods used by top-performing teams.

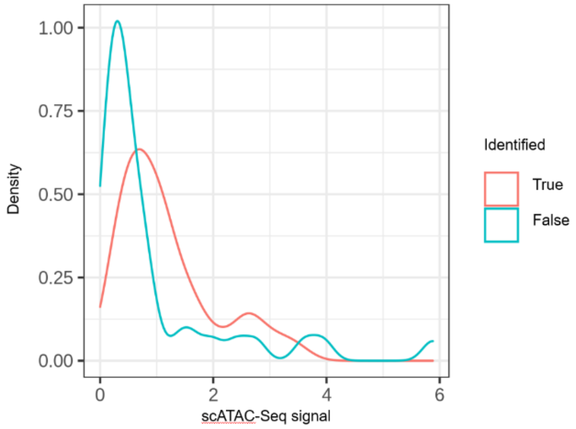


**Figure S4. ATAC-seq signal for peaks called or missed by teams.**
Density plots of ATAC-seq signal for enhancers identified by at least one team or missed by all teams.

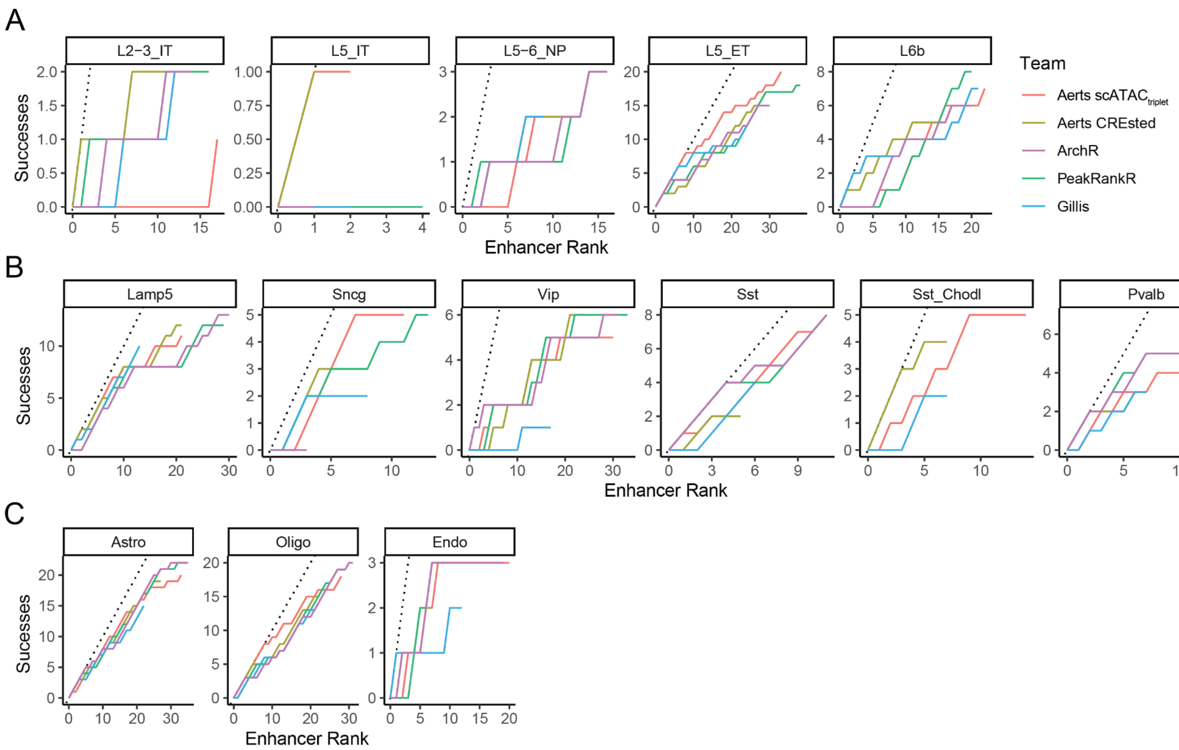


**Figure S5. Rates of functional enhancer identification vary by cell type and submission.**
Counts of On-Target enhancers identified for (A) excitatory and (B) inhibitory neuronal types and (C) non-neuronal types as a function of the number of validated enhancers for each type. Note that for some cell types (e.g. L5 IT), only some teams submitted any On-Target enhancer.

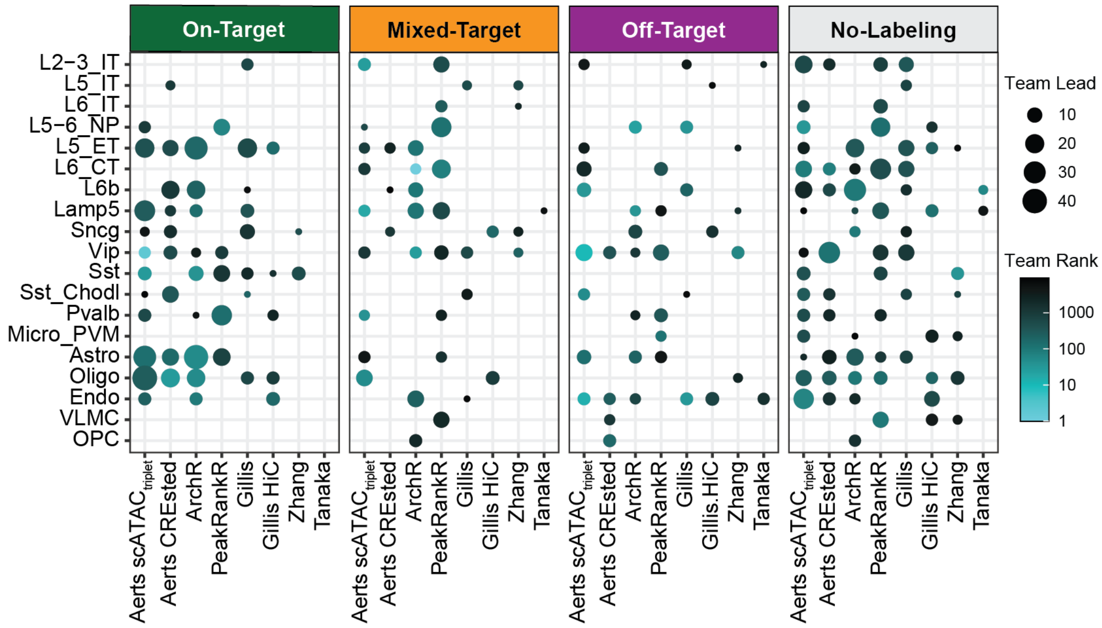

**Figure S6. Summary of the highest ranked enhancers across submissions.**

Enhancers were grouped by *in vivo* validated activity (e.g. On-Target), and point size indicates the number of enhancers that each submission scored the highest (i.e. lowest rank) compared to the other submissions. The color corresponds to the median rank of the enhancers included in each point. Note that better performing teams had larger, lighter points in the On-Target category (i.e., more enhancers that were scored higher) and smaller, darker points in the Mixed-Target, Off-Target, and No-Labeling categories (i.e., fewer enhancers that were scored lower).

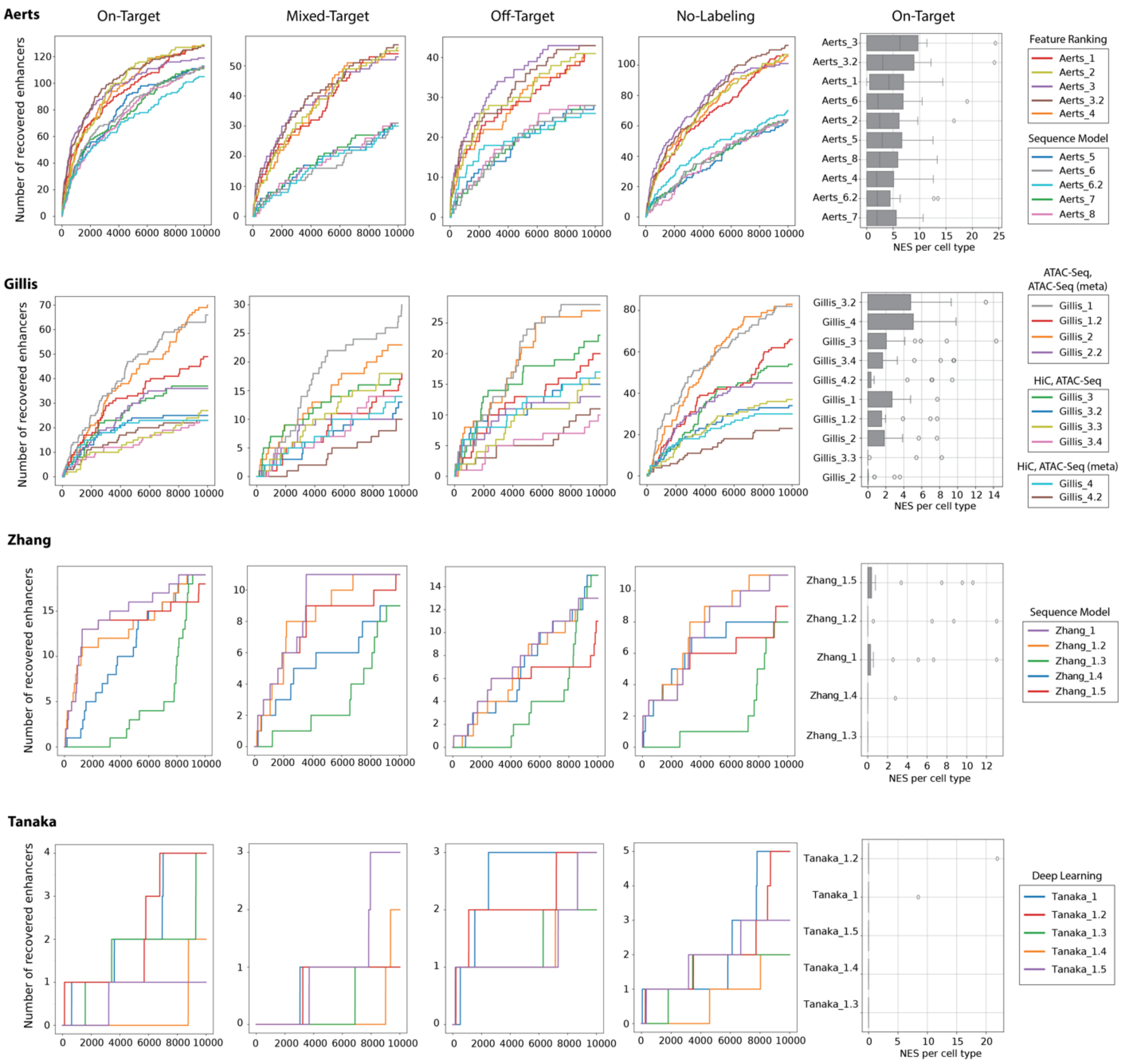


**Figure S7. Rates of enhancer identification and NES score for teams top submissions.**

Enhancers are grouped by *in vivo* specificity categories and each line indicates the number of enhancers recovered and at which rank at which a given enhancer was recovered by each submissions for teams. The normalized enrichment score (NES) is shown for each submission illustrating per cell type ranking. Teams submissions are grouped by method type to identify patterns in recovery curves.

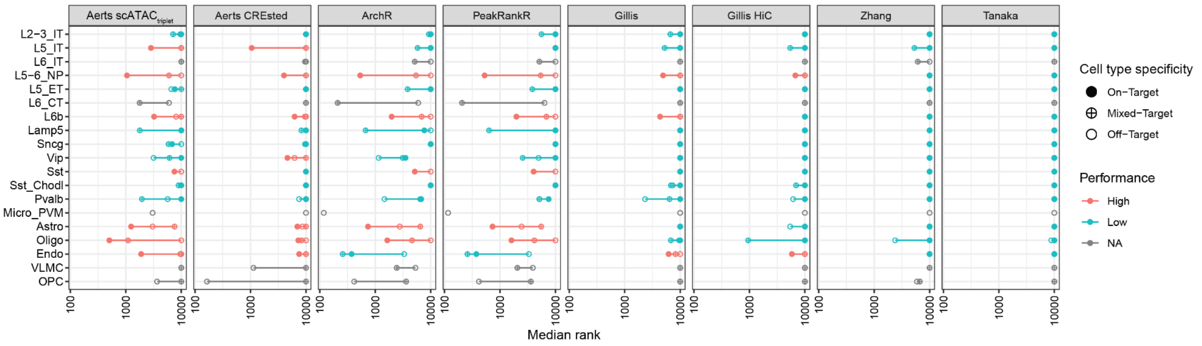


**Figure S8. Comparison of enhancer rankings across submissions, cell types, and *in vivo* labeling results.** Median ranks of enhancers grouped by submission, cell type, and specificity. Rankings were labeled “high performance” if the median rank of On-Target enhancers was higher than the rankings of Mixed-Target and Off-Target enhancers. If no On-Target enhancer rankings were submitted, then the performance was labeled “NA”.

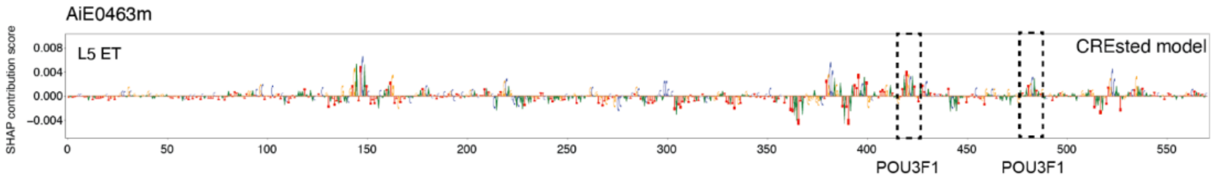


**Figure S9. *Pou3f1* enrichment in CREsted model for AiE0463m.**
The nucleotide contribution score track for enhancer AiE0463m for the L5 ET class of the Aerts CREsted model. Two *Pou3f1* motifs were identified around position 420 and 485.

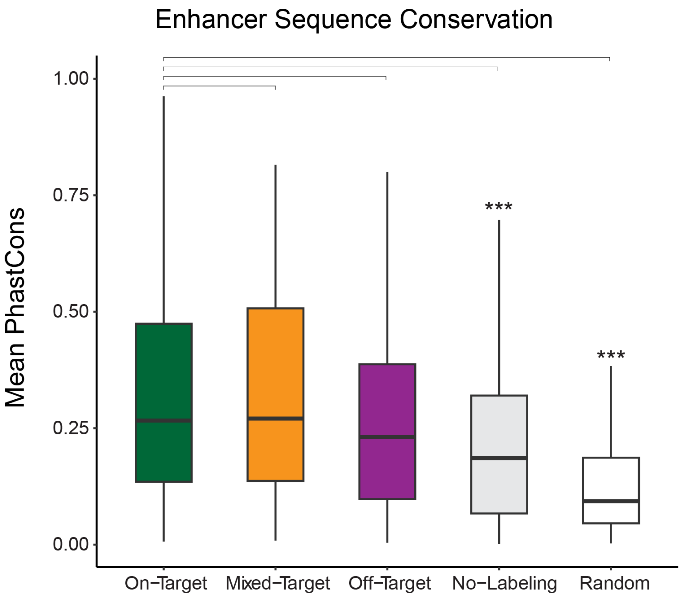


**Figure S10. DNA sequence conservation of screened enhancers grouped by cell type labeling results.** Comparison between On-Target and other enhancer categories. *** P < 0.0001, Wilcoxon rank-sum test two-sided, unpaired, Bonferroni-corrected P-values.

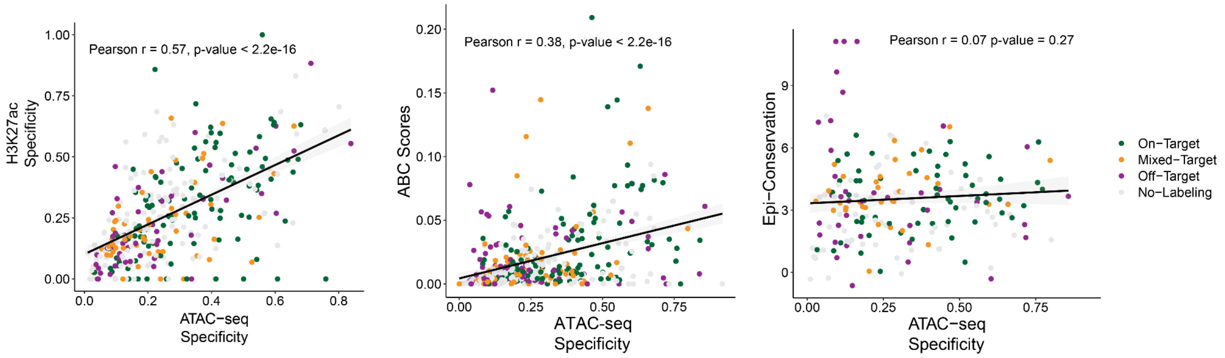


**Figure S11. Variable correlation of genomic features with ATAC-seq specificity.** H3K27ac specificity, ABC score, and open chromatin conservation between mouse and human compared to ATAC-seq specificity for all screened enhancers. Note that the candidate enhancers with the highest conservation of open chromatin (upper left quadrant of the third plot) were located at promoters and lacked cell type-specific activity.

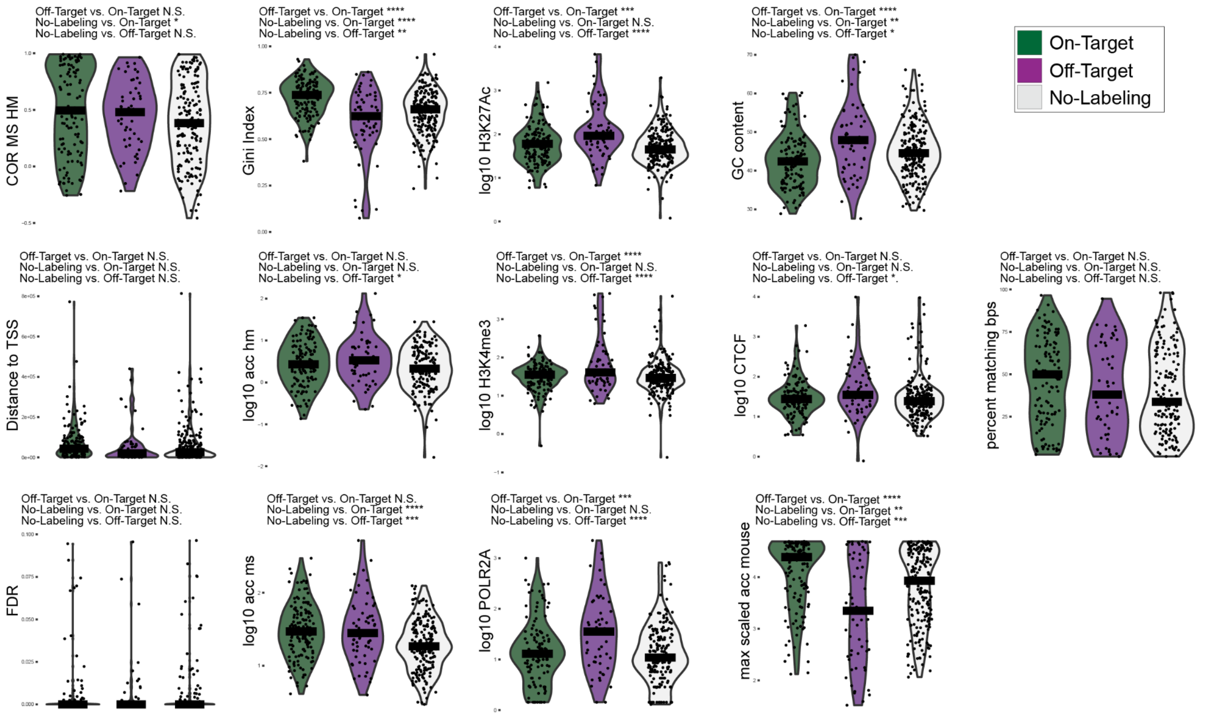


**Figure S12. Genomic features compared across high- and low-performance enhancers.**
Enhancer sequence features, species conservation and published bulk ChIP-seq data (ENCODE dataset ENCFF203KID) were used to train supervised random forest models to predict *in vivo* enhancer activity based on primary scoring data. Models were developed and optimized using 10-fold cross-validation in scikit-learn. Models were then tested using a held-out test set (70% training, 30% testing). Violin plots were generated for all features grouped by enhancer activity. * $P < 0.05$, ** $P < 0.01$, *** $P < 0.001$, **** $P < 0.0001$, ANOVA and Tukey post hoc tests, Bonferroni-corrected P-values.

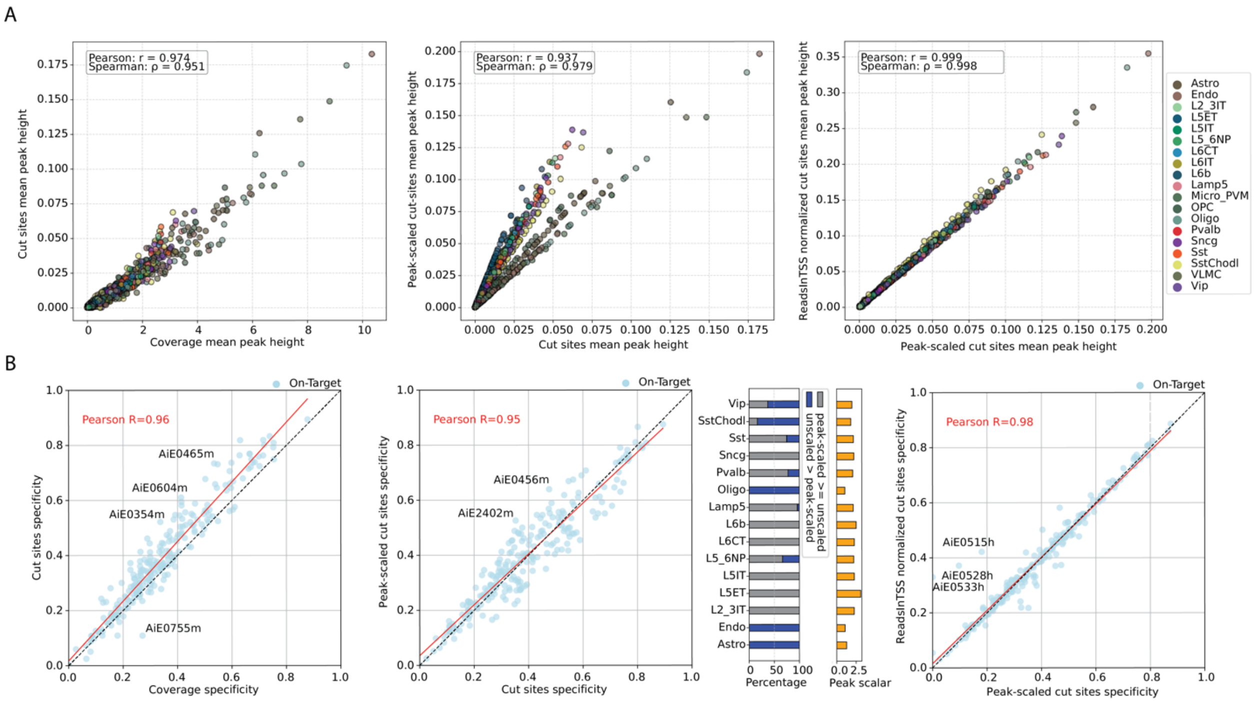


**Figure S13. Comparison of differential ATAC preprocessing methods.**

(**A**) Comparison of peak heights for all mouse enhancers after different preprocessing methods. (**B**) Scatter plot comparison of peak specificity for all on-target enhancers between different preprocessing methods. In the middle plot, the distributions of cell types for peaks which are more specific before and after peak scaling are given, together with the respective scalars.

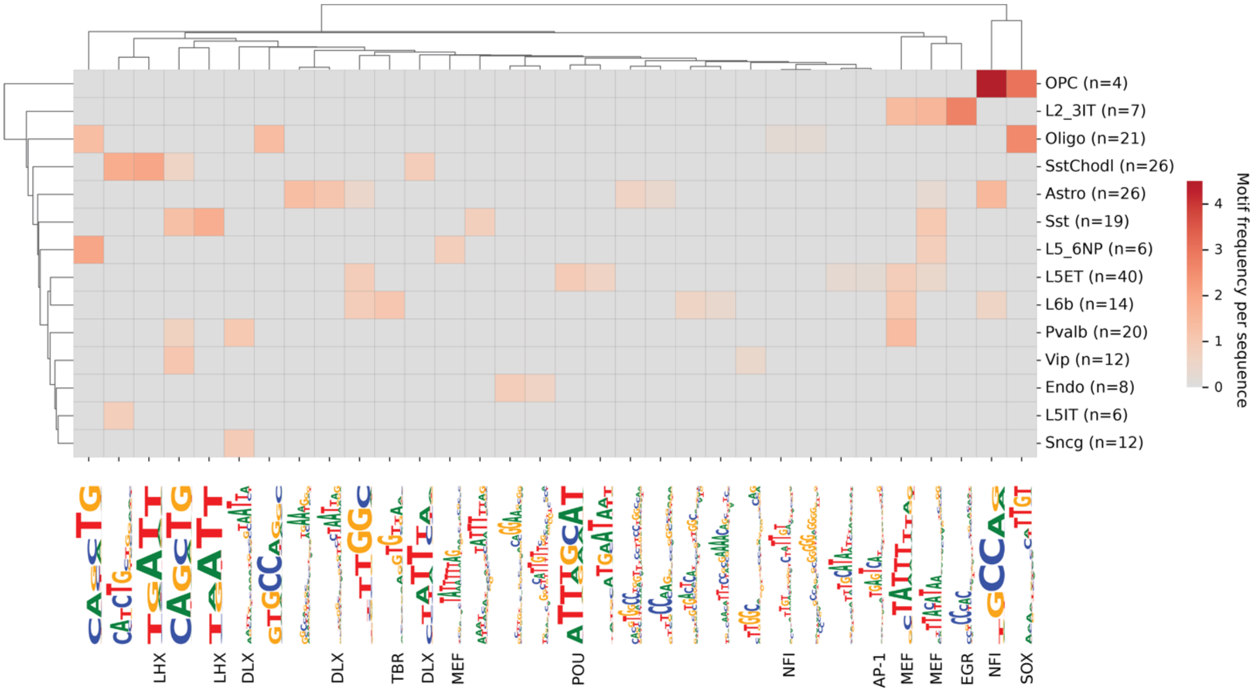


**Figure S14. Overview of transcription factor motifs within CREsted contribution scores for validated enhancers.**

Heatmap of consistently identified transcription factor motifs (columns) across each subclass (rows). The color intensity indicates the prevalence of a given motif in the respective subclass. Transcription factor motifs are named according to the associated motif family. Several sequence patterns do not correspond to known motifs and remain unlabeled.

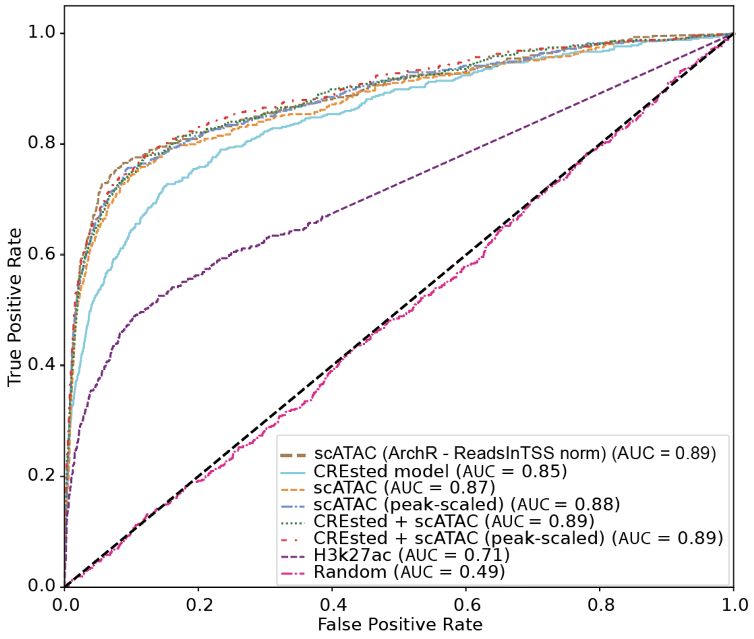


**Figure S15. Performance comparison of several models predicting *in vivo* enhancer activity.**
Area under the receiver operator curve to assess performance of enhancer ranking methods using the rescored enhancer activities.

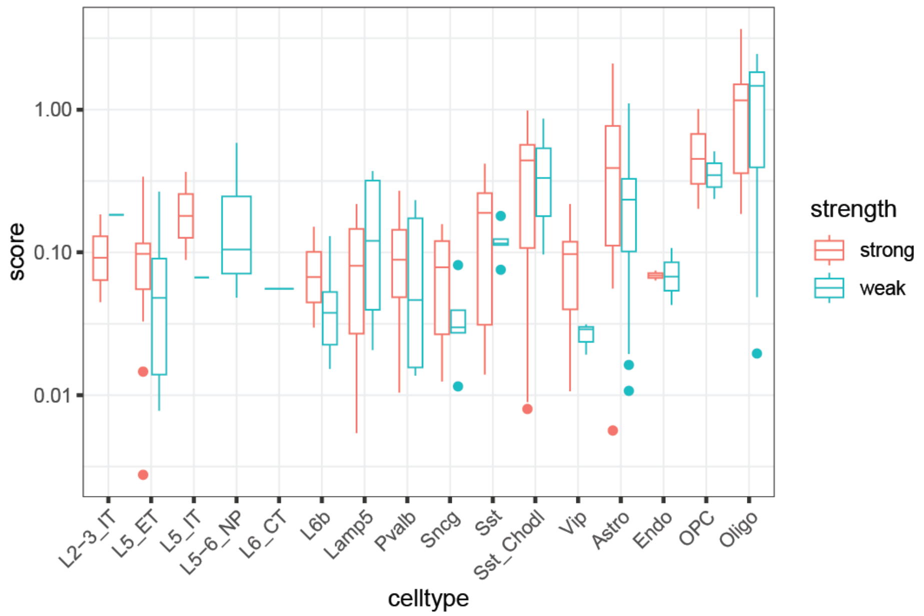


**Figure S16. Comparisons of DNA sequence model scores for strong and weak On-target enhancers.**
Boxplots of CREsted scores for candidate enhancers grouped by *in vivo* validated cell type target and strength of activity. Median +/- inter-quartile interval, outliers are points.

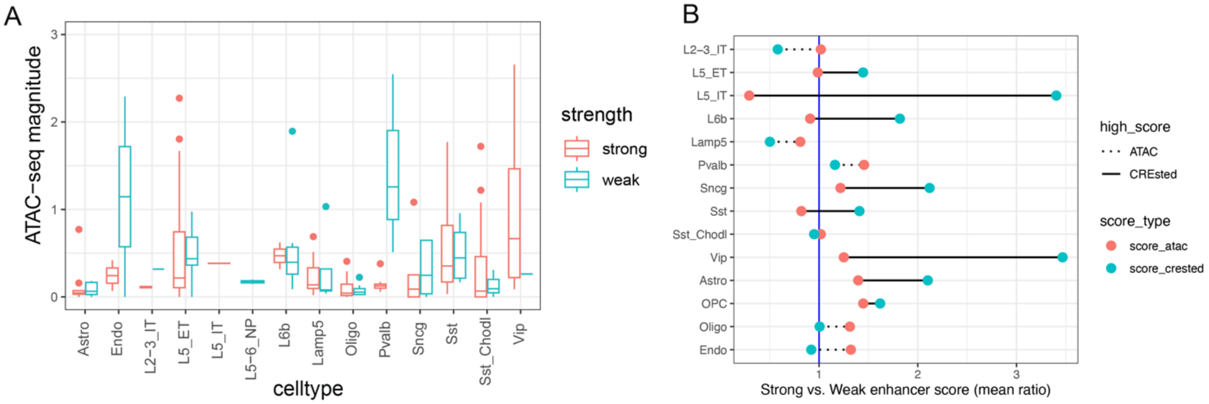


**Figure S17. Comparisons of ATAC-seq magnitude values for strong and weak On-target enhancers.**
**(A)** Boxplots of ATAC-seq magnitude values for candidate enhancers grouped by *in vivo* validated cell type target and strength of activity. Median +/- inter-quartile interval, outliers are points. **(B)** For each cell subclass, the average ATAC-seq magnitudes and CREsted sequence model scores were calculated for all enhancers, and the ratios for strong versus weak enhancers were plotted. CREsted (blue points) was better than ATAC-seq alone (red points) at predicting strong versus weak enhancers for the majority of subclasses, particularly for L5 IT and Vip neurons.

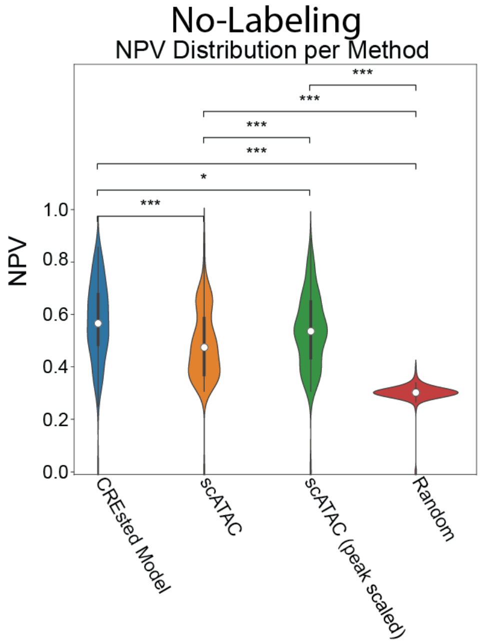


**Figure S18. CREsted model is better at identifying enhancers lacking *in vivo* activity.**
The negative predictive value (NPV) of the CREsted model is significantly higher than the NPVs of the initial and peak-scaled ATAC-seq models. * $P < 0.05$, *** $P < 0.001$, Wilcoxon rank-sum test two-sided, unpaired, Bonferroni-corrected P-values.

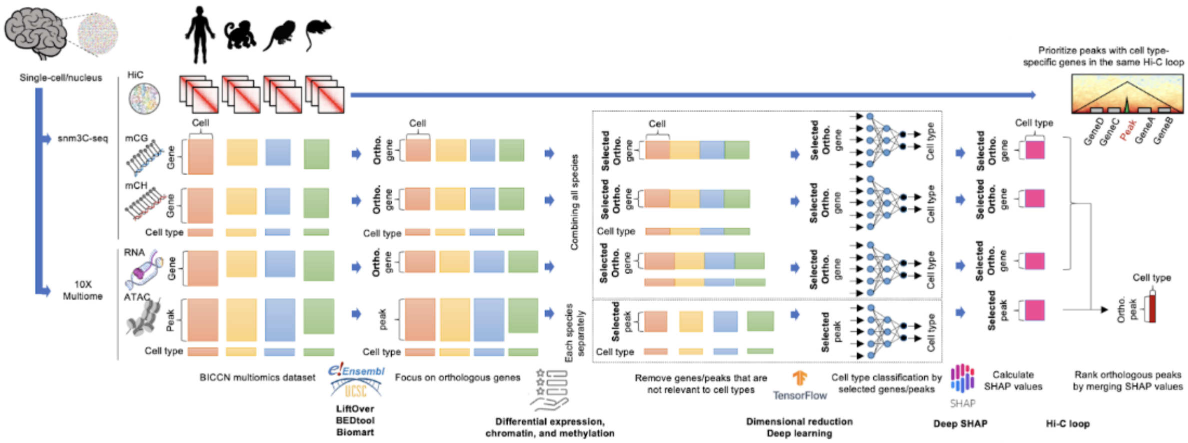


**Figure S19. Yoshiaki Tanaka lab enhancer prediction overview.** cisMultiDeep: Identifying Cell Type-Specific Cis-Regulatory Regions by Automatically-Tuned Deep Neural Network and SHAP.

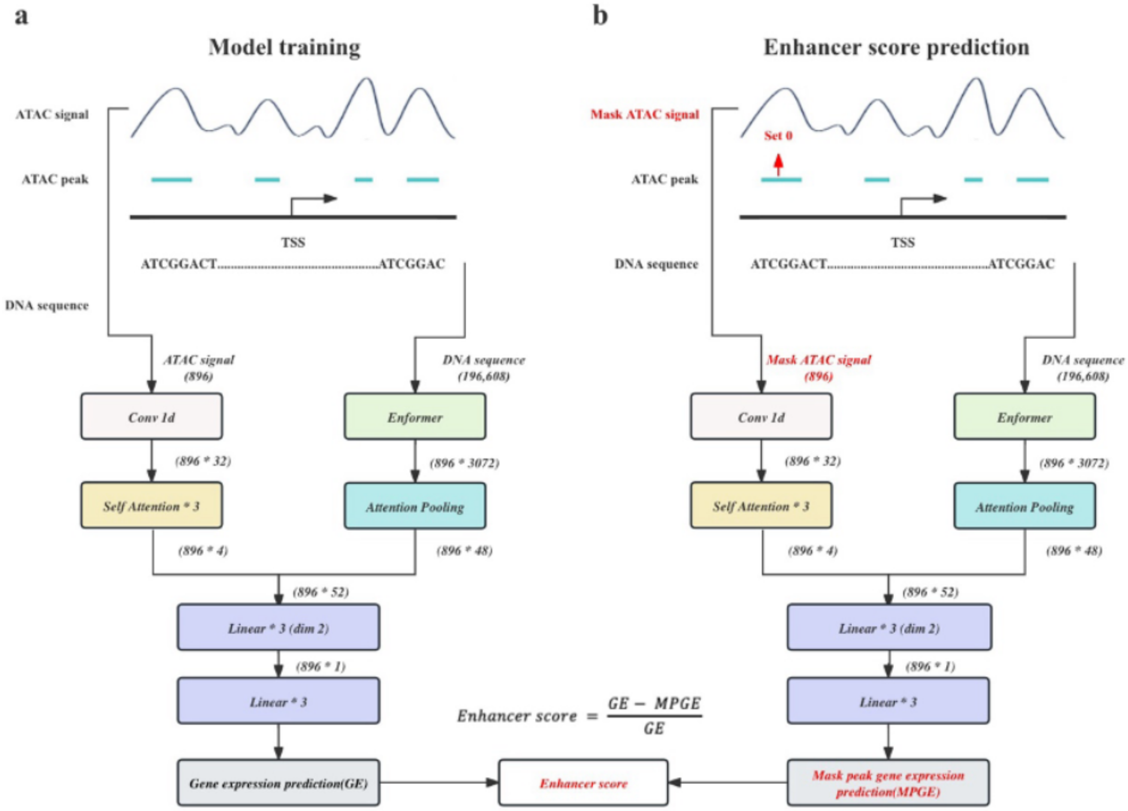


**Figure S20. Kai Zhang lab enhancer prediction overview.**

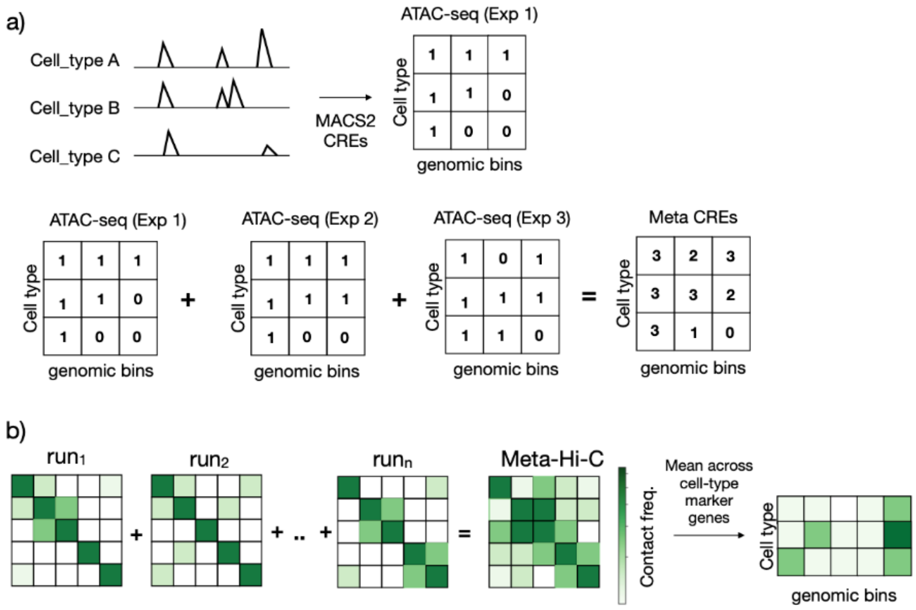


**Figure S21. Gillis lab enhancer prediction overview.**

***Supplemental Tables:***

**Supplement Table 1. Summary of validated enhancers.**
Table of validated enhancer data showing all counts.

**Supplement Table 2. Summary of team submissions.**
Table of benchmark metrics for all submissions and aliases used in manuscript figures.
